# Orchestrated Excitatory and Inhibitory Learning Rules Lead to the Unsupervised Emergence of Self-sustained and Inhibition-stabilized Dynamics

**DOI:** 10.1101/2020.12.30.424888

**Authors:** Saray Soldado-Magraner, Rodrigo Laje, Dean V. Buonomano

**Affiliations:** Department of Neurobiology, University of California, Los Angeles, CA, USA; Departamento de Ciencia y Tecnología, Universidad Nacional de Quilmes, Bernal, Argentina, and Consejo Nacional de Investigaciones Científicas y Técnicas (CONICET), Buenos Aires, Argentina; Department of Psychology University of California, Los Angeles, CA, USA

## Abstract

Self-sustaining neural activity maintained through local recurrent connections is of fundamental importance to cortical function. We show that Up-states—an example of self-sustained, inhibition-stabilized network dynamics—emerge in cortical circuits across three weeks of *ex vivo* development, establishing the presence of unsupervised learning rules capable of generating self-sustained dynamics. Previous computational models have established that four sets of weights (*W*_*E*←*E*_, *W*_*E*←*I*_, *W*_*I*←*E*_, *W*_*I*←*I*_) must interact in an orchestrated manner to produce Up-states, but have not addressed how a family of learning rules can operate in parallel at all four weight classes to generate self-sustained inhibition-stabilized dynamics. Using numerical and analytical methods we show that, in part due to the paradoxical effect, standard homeostatic rules are only stable in a narrow parameter regime. In contrast, we show that a family of biologically plausible learning rules based on “cross-homeostatic” plasticity robustly lead to the emergence of self-sustained, inhibition-stabilized dynamics.

## INTRODUCTION

Self-sustained patterns of neural activity maintained by local recurrent excitation underlie many cortical computations and dynamic regimes, including the persistent activity associated with working memory (Fuster and Jervey, 1981; Goldman-Rakic, 1995; Wang, 2001), asynchronous states (van Vreeswijk and Sompolinsky, 1998; Renart et al., 2010), and Up-states (Steriade et al., 1993; Timofeev et al., 2000). Recurrent excitation, however, also has the potential to drive pathological and epileptiform regimes (McCormick, 1989; Douglas et al., 1995; Steriade and Contreras, 1998). Converging theoretical and experimental evidence indicate that cortical circuits that generate self-sustained dynamics operate in an inhibition-stabilized regime, in which positive feedback is held in check by recurrent inhibition (Tsodyks et al., 1997; Brunel, 2000; Ozeki et al., 2009; Rubin et al., 2015; Rutishauser et al., 2015; Jercog et al., 2017; Sanzeni et al., 2020).

At the computational level self-sustained activity and inhibition-stabilized networks are often modeled as a simplified circuit composed of excitatory (*E*) and inhibitory (*I*) subpopulations of neurons with four classes of synaptic weights: *W*_*E*←*E*_, *W*_*E*←*I*_, *W*_*I*←*E*_, *W*_*I*←*I*_. Analytical and numerical analyses have shown that these weights must obey certain theoretically well-defined relationships in order to generate self-sustained, inhibition-stabilized dynamics (Tsodyks et al., 1997; Brunel, 2000; Ozeki et al., 2009; Rubin et al., 2015; Jercog et al., 2017). Yet, it is not known how the appropriate relationships between these four classes of weights could emerge in a self-organizing manner (Sadeh and Clopath, 2021). One possibility is that standard homeostatic forms of plasticity underlie the emergence of inhibition-stabilized networks. Homeostatic learning rules generally assume that excitatory weights are regulated in a manner proportional to the difference between some ontogenetically determined set-point and average neural activity (for both excitatory and inhibitory neurons)—and conversely that inhibitory weights onto excitatory neurons are regulated in the opposite direction (Turrigiano et al., 1998; van Rossum et al., 2000; Kilman et al., 2002; Turrigiano and Nelson, 2004; Peng et al., 2010). However, it remains an open question whether homeostatic rules can lead to the self-organized emergence of self-sustained, inhibition-stabilized networks.

At both the experimental and computational level one of the simplest and best-studied examples of self-sustained activity are Up-states (Steriade et al., 1993; Timofeev et al., 2000). Up-states are characterized by network-wide regimes in which excitatory and inhibitory neurons transiently shift from a quiescent Down-state to a depolarized state with low to moderate firing rates (Sanchez-Vives and McCormick, 2000; Neske et al., 2015; Bartram et al., 2017). Up-states occur spontaneously *in vivo* during anesthesia, slow-wave sleep, and quiet wakefulness (Steriade et al., 1993; Timofeev et al., 2000; Beltramo et al., 2013; Hromádka et al., 2013), in acute slices (Sanchez-Vives and McCormick, 2000; Shu et al., 2003; Fanselow and Connors, 2010; Sippy and Yuste, 2013; Xu et al., 2013; Sadovsky and MacLean, 2014; Neske et al., 2015; Bartram et al., 2017), and in organotypic cultures over the course of *ex vivo* development (Plenz and Kitai, 1998; Seamans et al., 2003; Johnson and Buonomano, 2007; Kroener et al., 2009; Motanis and Buonomano, 2015; Motanis and Buonomano, 2020). Furthermore, Up-state frequency appears to be homeostatically regulated—e.g., optogenetically stimulating cortical circuits over the course of days decreases Up-state frequency (Motanis and Buonomano, 2015; Motanis and Buonomano, 2020).

Consistent with previous results we first demonstrate that Up-states emerge in both excitatory and inhibitory neurons over the course of the first few weeks of *ex vivo* development, suggesting that local cortical circuits are programmed to homeostatically generate Up-states. We next used computational models and analytical methods to explore families of homeostatic learning rules that operate in parallel at all four synapse classes and lead to self-sustained, inhibition-stabilized dynamics. We show that when driving the network towards a self-sustained, inhibition-stabilized regime, standard forms of homeostatic plasticity are only stable in a narrow region of parameter space. This is in part a consequence of the paradoxical effect—in which an *increase* in excitatory drive to inhibitory neurons produces a net *decrease* in the firing rate of those same inhibitory neurons (Tsodyks et al., 1997; Ozeki et al., 2009; Rubin et al., 2015). We next developed a family of homeostatic learning rules that include “cross-homeostatic” influences, and lead to the unsupervised emergence of Up-states in the inhibition-stabilized regime in a robust manner. These rules are consistent with experimental data and generate explicit predictions regarding the effects of manipulations of excitatory and inhibitory neurons on synaptic plasticity.

## RESULTS

### Up-states emerge autonomously during ex vivo development

Up-states represent a transition from a quiescent state to a self-sustained network-wide dynamic regime in which both excitatory and inhibitory neurons are active (**Fig. 1A**). During Up-states the firing rate of excitatory neurons is relatively low (1-5 Hz) indicating that recurrent excitation is held in check by appropriately tuned inhibition (Neske et al., 2015; Jercog et al., 2017; Romero-Sosa et al., 2021). Computational studies have demonstrated that Up and Down states can be simulated as a bistable dynamical system composed of interconnected populations of excitatory (*E*) and inhibitory (*I*) neurons (**Fig. 1B**), in which Down-states represent a quiescent fixed point, and Up- or asynchronous states represent a second, non-trivial fixed-point attractor. In the Up regime recurrent excitation produces amplification, but the activity is held in check by rapid inhibition. The dynamics settles into a stable fixed-point attractor, and instantiates an example of an inhibition-stabilized network. The neural dynamics within two-population models is governed by four classes of synaptic weights *W*_*E*←*E*_, *W*_*E*←*I*_, *W*_*I*←*E*_, *W*_*I*←*I*_ (**Fig. 1B**, inset). Analytical and numerical studies have demonstrated that these four weights must obey certain “balanced” relationships in order to support the stable self-sustaining dynamics— e.g., if excitation is too strong, runaway (or saturated) excitation occurs, whereas if inhibition is too strong only the trivial quiescent fixed point will be stable (Tsodyks et al., 1997; van Vreeswijk and Sompolinsky, 1998; Brunel, 2000; Ozeki et al., 2009; Rubin et al., 2015; Jercog et al., 2017) (see **Section 2.2** in the Supplementary Material).

**Figure 1.**
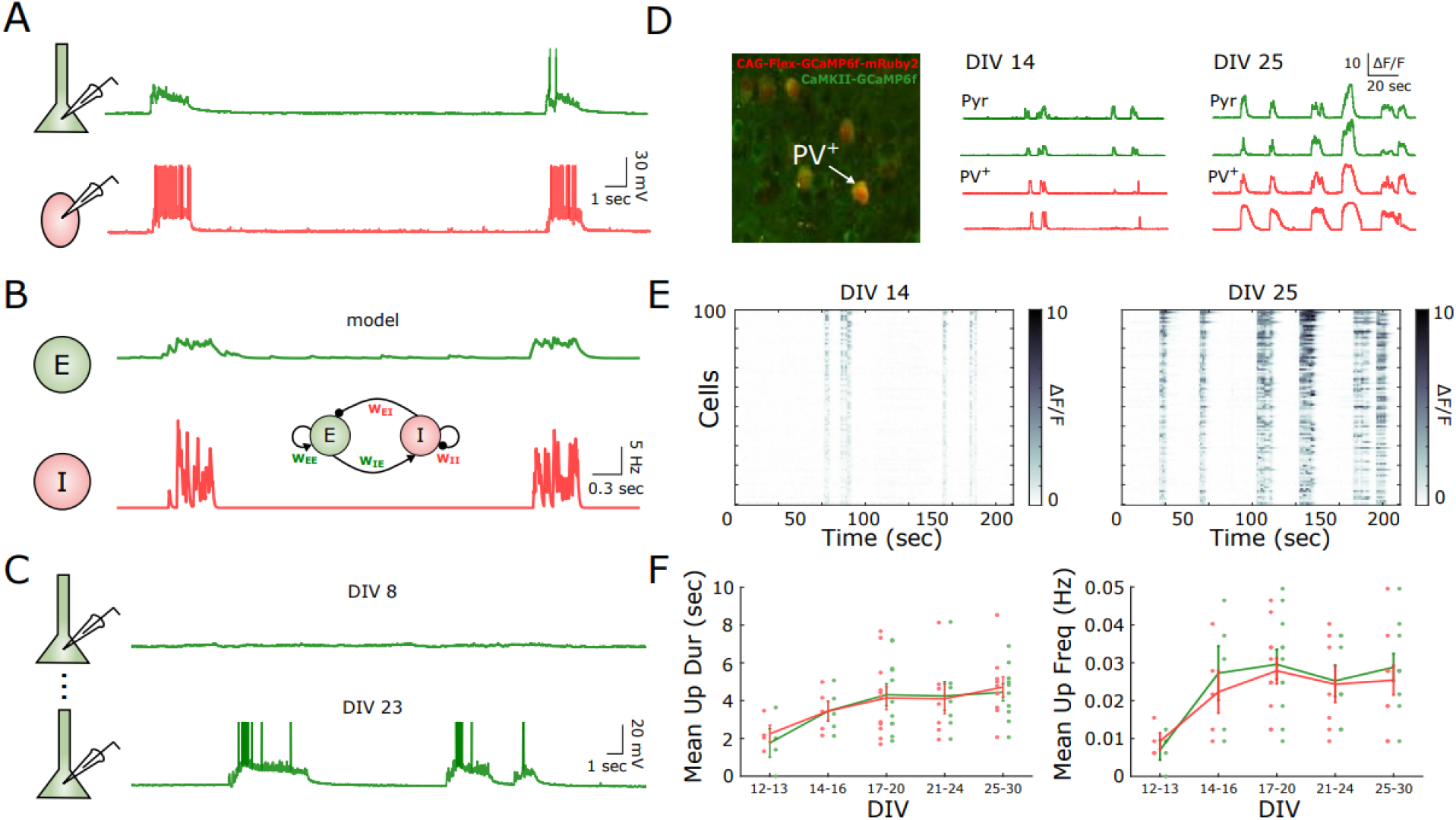
Up-states emerge autonomously over the course of *ex vivo* development. **(A)** Example of Up-states in simultaneously whole-cell recordings of a pyramidal (green) and parvalbumin (PV) positive inhibitory neuron (red). **(B)** Two-population firing rate model of Up-states. The schematic of the model is shown in the inset. The dynamics of the excitatory (green) and inhibitory (red) populations are governed by four synaptic weights, *W*_*E*←*E*_, *W*_*E*←*I*_, *W*_*I*←*E*_, and *W*_*I*←*I*_. Traces correspond to the firing rate of each of the populations in the presence of external noise. **(C)** Spontaneous activity recording of a pyramidal neuron at 8 and 23 days *in vitro* development (DIV). Up-states are present only at later developmental stages. **(D)** Two-photon calcium imaging recording of excitatory and PV^+^ neurons at different stages of development. Organotypic slices of Cre-PV mice were transfected with pAAV-CAG-Flex-GCamp6f-mRuby2 and pAAV-CaMKII-GCamp6f. Image shows an example slice with a PV^+^ neuron expressing both, GCamp6f and the mRuby2 red marker. Traces show the spontaneous calcium activity of 2 example PV^+^ and excitatory cells at 14 and 25 DIV. Up-states can be observed more prominently at later stages (see Methods for the definition and quantification of Up-states). **(E)** Spontaneous calcium activity of 100 example cells at 14 and 25 DIV. Synchronous activity events correspond to Up-states. **(F)** Evolution of the mean Up-state duration and frequency over the course of *ex vivo* development for excitatory (green) and PV^+^ neurons (red). A significant increase in mean Up duration (Two-way ANOVA: F_4,64_ = 3.54, p = 0.011) and frequency (Two-way ANOVA: F_4,64_ = 4.75, p = 0.002) was observed over developmental time, with no statistical effect of neuron type (F_4,64_ = 0.13, p = 0.97) or interaction effect (F_4,64_ = 0.09, p = 0.98).

In most computational models the set of four weights is determined analytically or through numerical searches. In contrast, recordings in cortical organotypic cultures show that Up-states autonomously develop over the course of *ex vivo* development (Plenz and Kitai, 1998; Seamans et al., 2003; Johnson and Buonomano, 2007; Kroener et al., 2009; Motanis and Buonomano, 2015; Motanis and Buonomano, 2020). Early in development, at 8 days-in-vitro (DIV-8) most of the neurons are silent, while at later stages (DIV-23) spontaneous Up-states are observed (**Fig. 1C**). Here we further characterized the emergence of Up-states and asked whether the development of activity is in sync for both excitatory and inhibitory neurons. Using two-photon calcium imaging, we recorded the spontaneous activity in excitatory neurons and PV^+^-inhibitory neurons by expressing GCamp6f under the CaMKII and Flex promoters in organotypic cultures of PV-Cre mice. Calcium imaging at DIV 12-13 revealed infrequent and short bouts of synchronous activity. By DIV 14-16 Up-states were observed, and over the entire four-weeks of *ex vivo* development there was an increase and stabilization of Up-state frequency and duration in both excitatory and inhibitory neurons (**Fig. 1D-F**)—suggesting the Up-states emerge in a co-dependent manner in both populations.

The observation that Up-states emerge autonomously during *ex vivo* development indicates that synaptic learning rules are in place to orchestrate the unsupervised emergence of Up-states. Since Up-states emerge autonomously over the course of development in *ex vivo* cortical networks, and because all four weight classes have been observed to undergo synaptic plasticity in experimental studies, we next asked how the stable self-sustained dynamics characteristic of Up-states might emerge in a self-organizing manner.

### Standard homeostatic learning can only account for stable self-sustained activity in a narrow parameter regime

One attractive possibility is that cortical neurons are homeostatically programmed to generate Up-states. Specifically, that both excitatory and inhibitory neurons exhibit ontogenetically programmed firing rate setpoints during Up-states, and they homeostatically adjust their excitatory and inhibitory weights to reach these target setpoints. Homeostatic learning rules are traditionally defined by changes in synaptic weights that are proportional to an “error term” defined by the difference between the setpoint and the neurons average activity levels (Turrigiano et al., 1998; van Rossum et al., 2000; Kilman et al., 2002; Turrigiano and Nelson, 2004; Liu and Buonomano, 2009; Peng et al., 2010; Vogels et al., 2011), e.g., Δ*W*_*E*←*E*_ ∝ *E*_*set –*_ *E*_*avg*_ where any departure of the excitatory activity *E*_*avg*_ from the setpoint *E*_*set*_ would lead to a compensatory correction in the value of the weight *W*_*E*←*E*_.

We first asked is stable self-sustained dynamics can emerge in a standard two-population model (Jercog et al., 2017; see Methods) through homeostatic mechanisms. We initialized the four weights (*W*_*E*←*E*_, *W*_*E*←*I*_, *W*_*I*←*E*_, *W*_*I*←*I*_) of the model at random values and applied a standard family of homeostatic learning rules to all four weights classes (**Fig. 2A**). It is well established that PV^+^-inhibitory neurons have higher firing rates than pyramidal neurons during Up-states (Neske et al., 2015; Romero-Sosa et al., 2021), thus based on experimental data we set the setpoints for the *E* and *I* populations during Up-states to 5 and 14 Hz, respectively (Romero-Sosa et al., 2021). We first asked whether the set of four standard homeostatic learning rules can lead to stable self-sustained dynamic regime (representing a permanent Up-state) in response to a brief external input (low levels of noise were used to avoid spontaneous Up↔Down transitions).

**Figure 2.**
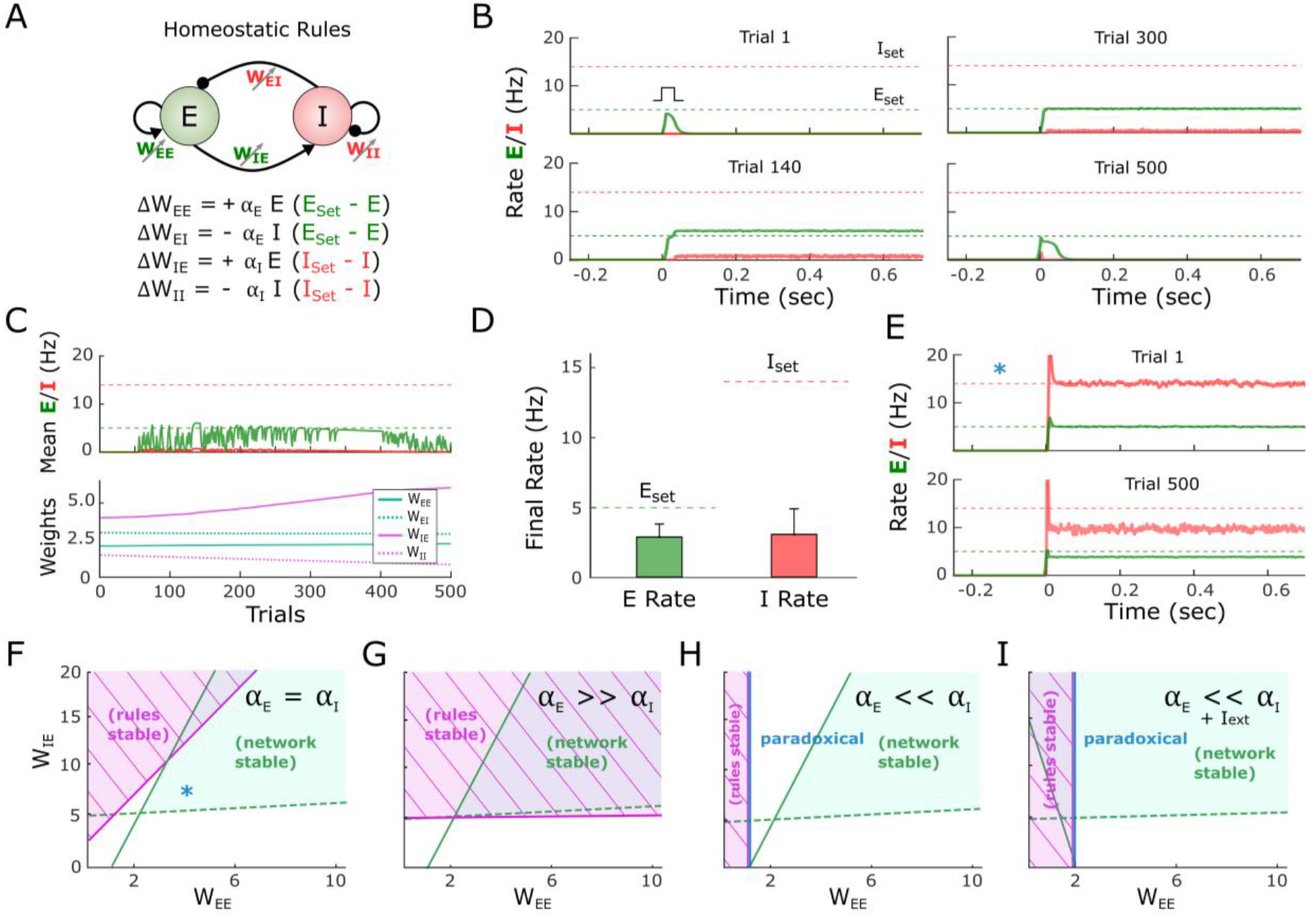
Standard homeostatic rules are only stable in a narrow parameter regime. **(A)** Schematic (top) of the population rate model in which the four weights are governed by a family of homeostatic learning rules (bottom). **(B)** Example simulation of the network over the course of simulated development. Each plot shows the firing rate of the excitatory and inhibitory population over the course of a trial in response to a brief external input. *E*_*set*_*=5* and *I*_*set*_=14 represent the target homeostatic setpoints. Weights were initialized to *W*_*EE*_=2.1, *W*_*EI*_=3, *W*_*IE*_=4, and *W*_*II*_=2. Note that while an evoked Up-state emerges by Trial 200 the firing rates do not converge to their setpoints, and by Trial 500 the Up-state is no longer observed. **(C)** Average rate across trials (upper plot) for the excitatory and inhibitory populations for the data shown in (B). Weight dynamics (bottom plot) produced by the homeostatic rules across trials for the data shown in (B). **(D)** Average final rate for 100 independent simulations with different weight initializations. Data represents mean ± SEM. **(E)** Simulation of a network starting with weights that generate Up-states that match the *E*_*set*_=5 and *I*_*set*_=14 Hz setpoints (Trial 1, top). After 500 trials the network has diverged from its setpoints, indicating the synaptic learning rules are unstable. Weights were initialized to *W*_*EE*_=5 *W*_*EI*_=1.09 *W*_*IE*_=10 *W*_*II*_=1.54. **(F)** Analytical stability regions of the neural and learning rule subsystems as a function of the free weights *W*_*EE*_ and *W*_*IE*_. (note that once a *W*_*EE*_ and *W*_*IE*_ are set to generate an Up-state with specific *E*_*set*_ and *I*_*set*_ values, *W*_*EI*_ and *W*_*II*_ are fully determined by *W*_*EE*_ and *W*_*IE*_, respectively). Here the stability plot is obtained by considering equal learning rates for all four learning rules (as used for panels B-E). Blue asterisk corresponds to the initial conditions shown in Panel D (top). **(G)** Similar to F but with *α*_*E*_ >> *α*_*I*_. **(H)** Similar to F but with but with *α*_*E*_ << *α*_*I*_. To the right of the blue line, the network is in a paradoxical regime (defined by the condition W_EE*_g_E_ – 1 > 0) **(I)** Condition of stability of the neural system and learning rule system when the learning rate on the inhibitory neuron dominates and an external excitatory current is applied to the excitatory neuron. The current produces an enlargement of the stability region of the neural subsystem. Right of blue line shows the area where the network is in a paradoxical regime.

Although the rules are homeostatic in nature (e.g., if *I* is below *I*_*set*,_ an increase of *W*_*I*←*E*_ and a decrease in *W*_*I*←*I*_ would be induced), in the example shown in **Fig. 2B-C** the network failed to converge to a stable Up-state (**Fig. 2B-C**). Initially (Trial 1) an external input to the excitatory population does not engage recurrent activity because *W*_*E*←*E*_ is too weak. By Trial 200 the weights have evolved and the brief external input triggers an Up-state, but the activities *E* and *I* do not match the corresponding setpoints—the network is in a nonbiologically observed regime in which *E* > *I*—so the weights keep evolving. By Trial 600 *E* = *E*_*set*_ but *I* < *I*_*set*_, and rather than converging to *I*_*set*_, the network returns to a regime without an Up-state by Trial 1000. At that point both setpoint error terms have increased, leading to continued weight changes (**Fig. 2C)**. Results across 100 simulations with different weight initializations (see Methods) further indicate that the standard homeostatic rules are ineffective at driving *E* and *I* towards their respective setpoints and generating stable self-sustained dynamics (**Fig. 2D**).

To gain insights into why a family of homeostatic learning rules that might intuitively converge fails to do so, we can consider the case in which a network is initialized to a set of weights that already match *E*_*set*_ and *I*_*set*_ (**Fig. 2E**). Although the neural subsystem alone is stable at this condition (Trial 1), small fluctuations in *E* and *I* cause the homeostatic rules to drive the weight values and the average activity of the network away from the setpoints (Trial 500). It is possible to understand this instability by performing an analytical stability analysis. Specifically, a two-population network in which the weights undergo plasticity can be characterized as a dynamical system composed of two subsystems: the neural subsystem composed of the two differential equations that define *E* and *I* dynamics, and the synaptic learning rule subsystem defined by the four learning rules (see **Section 2.1** in the Supplementary Material). We make use of the two very different time scales of the neural (fast) and learning rule (slow) subsystems to perform a quasi-steady state approximation of the neural subsystem; then we compute the eigenvalues of the four-dimensional learning rule subsystem, and finally get an analytical expression for the stability condition of the learning rules (see **Section 2.3** in the Supplementary Material). For the entire system to be stable, both the neural and learning rules subsystems have to be stable. For the results presented in **Fig. 2B-E** we assumed the learning rates driving plasticity onto the excitatory (*α*_*E*_) and inhibitory neurons (*α*_*I*_) to be equal. Under these conditions, the standard homeostatic rules are mostly unstable for biologically meaningful parameter values in which the neural system is stable. An example of this result is shown in **Fig. 2F** for a particular set of parameter values. Critically, **Fig. 2F** shows that the stability region of the neural subsystem, i.e., an inhibition-stabilized network (Ozeki et al., 2009; Jercog et al., 2017), is almost entirely within the region where the homeostatic learning rule system is unstable. Only when plasticity onto the excitatory neuron is significantly faster (*α*_*E*_ >>*α*_*I*_) is there a substantial region of overlap between the stability of the neural and learning rules subsystems (**Fig. 2G**, see Supplementary Material, **Section 1.1**).

Because inhibitory neurons seem to undergo homeostatic plasticity as quickly or more quickly than excitatory neurons (Keck et al., 2011; Kuhlman et al., 2013; Gainey et al., 2018; Ma et al., 2019) we conclude that standard homeostatic rules by themselves do not account for the emergence of stable self-sustained and inhibition-stabilized dynamics. Similarly, a combination of analytical and numerical methods also indicates that variants of these homeostatic rules, such as synaptic scaling (Turrigiano et al., 1998; van Rossum et al., 2000; Sullivan and de Sa, 2006) are also only stable in a narrow region of parameter space (see Supplementary Material, **Section 1.5**). We next show that the inherent instability of standard homeostatic learning rules is related to the paradoxical effect.

### The paradoxical effect hampers the ability of homeostatic rules to lead to self-sustained activity

The inability of the homeostatic learning rules to generate stable Up-states is in part a consequence of the paradoxical effect, a counterintuitive, yet well described, property of two-population models of Up-states and inhibition-stabilized networks (Tsodyks et al., 1997; Ozeki et al., 2009). Specifically, if during an Up-state one increases the excitatory drive to the inhibitory population, the net result is a decrease in the firing rate of the inhibitory units. This paradoxical effect can be understood in terms of the *I*→*E*→*I* loop: the increased inhibitory drive leads to a lower steady-state rate for *E*, but this new steady-state value requires a decrease in the *I* firing rate to maintain an appropriate E/I balance (in effect, the decrease in *E* decreases the drive to *I* by more than the external increase to *I*). This paradoxical effect has profound consequences for learning rules that attempt to drive excitatory and inhibitory weights to their setpoints.

The relationship of the paradoxical effect and the homeostatic rules performance is presented in **Fig. 2H**. The region of stability for the homeostatic learning rules is shown in a parameter regime where inhibitory plasticity is much faster (α_E_ << α_I_). Contrary to when excitatory plasticity dominates, the region of stability is small, and there is no overlap with the region of stability of the neural subsystem. Crucially, the boundary of the stability region of the learning rule coincides with the condition for the paradoxical effect to be present (right of the blue line in **Fig.2H**, see Supplementary Material, **Sections 2.2 and 2.3**). Under these conditions, the rules can only be stable when the network is not in an inhibition-stabilized regime. If a network regime with non-zero *E* would be forced to exist in that region (for example, via tonic external current, **Fig. 2I**), it would only be stable in the non-paradoxical region with the learning rules in place (see **Section 2.5** in the Supplementary Material).

To understand the impact of the paradoxical effect on homeostatic learning rules consider a network state in which the *I* rate falls significantly below its setpoint, and the *E* rate is close to its setpoint (**Fig. 3A**). In order to reach the *I* setpoint, homeostatic plasticity in the inhibitory neuron would intuitively result in an increase of *W*_*I*←*E*_. However, because of the paradoxical effect an increase in *W*_*I*←*E*_ actually makes *I* decrease (**Fig. 3B**)—thus increasing the error term *I*_*set*_ – *I*. To increase the steady-state inhibitory rate we can “anti-homeostatically” decrease the excitatory weight onto the inhibitory neurons (**Fig. 3C)**. This simple example shows the complexity of designing a coherent set of rules in such a coupled system (see an analysis of the paradoxical effect in **Section 2.2** of the Supplementary Material). This analysis also explains why homeostatic learning rules can lead to self-sustained activity at the appropriated setpoints when *α*_*E*_ >>*α*_*I*_. Essentially by allowing plasticity onto the *E* population to be faster one overcomes the counterproductive homeostatic plasticity associated with the paradoxical effect.

**Figure 3.**
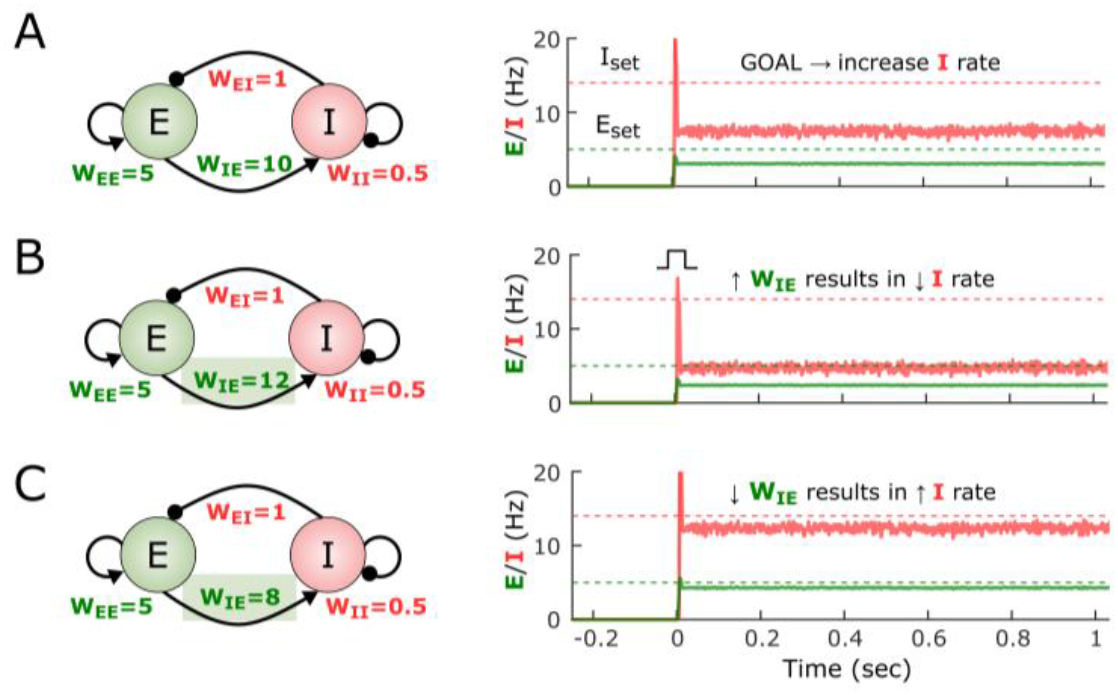
The paradoxical effect constrains the learning rules that can lead to Up-states. **(A)** Example of the self-sustained dynamics of a two-population model with weight values shown in the diagram. Both the *E* and *I* firing rates fall below their respective setpoints. The objective is to adjust the weights so that the *E* and *I* activity match their setpoints. **(B)** An increase of *W*_*IE*_ from 10 to 12 results in a paradoxical decrease of the *I* rate. **(C)** Because of the paradoxical effect an effective way to increase the steady-state *I* firing rate is to decrease its excitatory drive (i.e., *W*_*IE*_).

The interaction between the paradoxical effect and homeostatic plasticity in inhibitory neurons leads to the question of whether anti-homeostatic plasticity rules may be more effective that standard homeostatic rules—e.g., Δ*W*_*I* ← *E*_ ∝ -(*I*_*set*_ - *I*_*avg*_). Thus, we also examined a number of hybrid families of learning rules with different combinations of homeostatic and anti-homeostatic rules. Indeed, some hybrid families exhibited large degrees of overlap between the stable regions of the network and learning rules subsystems. However, numerical simulations revealed that these rules were mostly ineffective in driving networks to self-sustained activity at the target setpoints (Supplemental Material, **Section 1.2**, and Supplementary **Fig.S1**). These two results are not inconsistent because the stability analysis speaks to cases when the network is initialized to weights that satisfy *E*_*set*_ and *I*_*set*_, not whether the rules will drive network activity into these stable areas from any initial state including a pre-developmental state. Thus, we interpret these results as meaning that while anti-homeostatic plasticity can contribute to stability of this dual dynamical system, anti-homeostatic plasticity is ineffective at driving the dynamics towards setpoints (in other words, that anti-homeostatic plasticity might allow for stable Up-state but does not necessarily generate sizable basins of attraction around Up-states).

### A novel cross-homeostatic rule robustly leads to the emergence of self-sustained Up-states

Given that a standard set of homeostatic learning rules did not robustly lead to self-sustained dynamics we explored alternative learning rules. By defining a loss function based on the sum of the excitatory and inhibitory errors we analytically derived a set of learning rules using gradient descent (see **Section 3** in the Supplementary Material). This approach led to mathematically complex and biologically implausible rules; however, approximations and simulations inspired a simple class of learning rules that we will refer to as cross-homeostatic (see Methods). The main characteristic of this set of rules is that the homeostatic setpoints are “crossed” (**Fig. 4A)**. Specifically, the weights onto the excitatory neuron (*W*_*E*←*E*_ and *W*_*E*←*I*_) are updated to minimize the inhibitory error while weights into the inhibitory neuron (*W*_*I*←*E*_ and *W*_*I*←*I*_) change to minimize the excitatory error. Although apparently non-local, from the perspective of an excitatory neuron these rules can be interpreted as cells having a setpoints for the inhibitory input current onto the cell. Such input could be read by a cell as the activation of metabotropic receptors (e.g, GABA_b_ and mGlu; see Discussion). Indeed, a similar cross-homeostatic rule has been derived for *W*_*I*←*E*_ weights (Mackwood et al., 2021).

**Figure 4.**
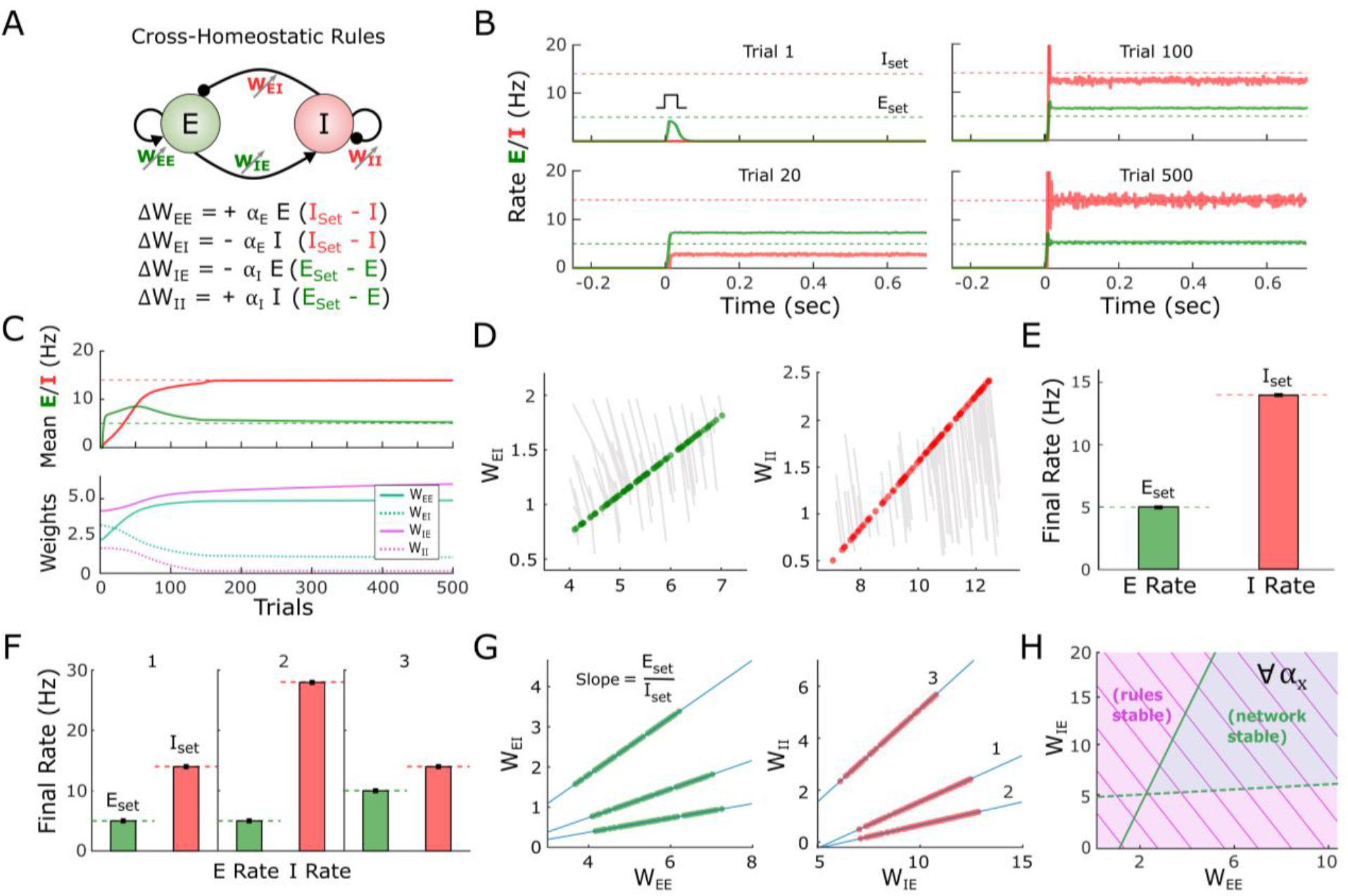
A family of cross-homeostatic learning rules robustly lead to self-sustained dynamics at *E*_*set*_ and *I*_*set*_. **(A)** Schematic of the network model and the family of cross-homeostatic learning rules. **(B)** Example network dynamics across simulated development. The network is initialized with weights that do not lead to self-sustained dynamics in response to an external input (Trial 1, weights are initialized to *W*_*EE*_=2.1 *W*_*EI*_=3 *W*_*IE*_=4 *W*_*II*_=2). By Trial 20 a stable Up-state is observed, but at firing rates far from the target setpoints (dashed lines). By Trial 500 the network has converged to an Up-state in which *E* and *I* firing rate match their respective setpoints **(C)** Average rate across trials (upper plot) for the excitatory and inhibitory populations for the data shown in (B). Weight dynamics (bottom plot) induced by the cross-homeostatic rules across trials for the data shown in (B) **(D)** Weight changes for 100 different simulations with random weight initializations (see Methods). Lines show change from initial to final (circles) weight values. **(E)** Average final rates for 100 independent simulations with different weight initializations shown in (D). Data represents mean ± SEM. **(F)** Final rates for the excitatory and inhibitory subpopulations after learning with same starting conditions as in (D) and (E) but for different setpoints. 1: *E*_*set*_=5, *I*_*set*_=14; 2: *E*_*set*_=5, *I*_*set*_=28; 3: *E*_*set*_=10, *I*_*set*_=14. Data shown in (D) and (E) corresponds to 1. Data represents mean ± SEM. **(G)** Final weight values for homeostatic plasticity simulations for the three different pairs of setpoints shown in (F). Blue lines correspond to the theoretical linear relationship between the excitatory and inhibitory weights at a fixed-point obeying *E*_*set*_ and *I*_*set*_. The slope of the line is defined by the ratio of the setpoints (see Methods). **(H)** Analytical stability regions of the neural subsystem and learning rule subsystem as a function of *W*_*EE*_ and *W*_*IE*_. The stability condition holds for any possible combination of learning rates (see **Section 2.4** in the Supplementary Material)

An example of the performance of the cross-homeostatic rules is shown in **Fig. 4B-C**. After an initial phase with no self-sustained firing (Trial 1), recurrent activity reaches a stable Up-state (Trial 20), whose average rate continues to converge towards its defined setpoints (Trial 100) until the learning rule system reaches steady state (Trial 500). The average *E* and *I* rates of the network evolve asymptotically towards the defined setpoints, as the weights evolve and converge (**Fig. 4C)**. Across different weight initializations the rules proved effective in driving the mean Up-state activity of the network to the target *E* and *I* setpoints, and led to balanced dynamics (**Fig. 4D-E**). The weight trajectory from its initial value to its final one is shown for 100 different simulations (**Fig. 4D**). Each line corresponds to individual experiments with different initializations. Circles indicate the final values of the weights. Independently of the initial conditions, the weights converge to a line attractor (actually a 2D plane attractor in 4D weight space; see **Section 2.1** in the Supplementary Material). Note that this attractor refers to the sets of weights that generate Up-states where *E* and *I* activity matches *E*_*set*_ and *I*_*set*_ respectively. That is, for a given pair of setpoints (*E*_*set*_, *I*_*set*_) the final values of the weights *W*_*E*←*I*_ and *W*_*I*←*I*_ are linear functions of the “free” weights *W*_*E*←*E*_ and *W*_*I*←*E*_, respectively. This is a direct consequence of the steady state conditions for the nontrivial fixed-point of the two-population model (Tsodyks et al., 1997; Ozeki et al., 2009; Jercog et al., 2017), where the slope of the line is defined by the setpoints *E*_*set*_/*I*_*set*_ (see Methods). For example, to satisfy 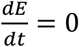 in the Up-state fixed point, the net excitation and inhibition must obey a specific “balance”, meaning that once *W*_*E*←*E*_ or *W*_*E*←*I*_ is determined, the other is analytically constrained for a given set of setpoints and parameters. Once the weights reach this specific relationship, the *E* and *I* rates reach their corresponding *E*_*set*_ and *I*_*set*_ values (**Fig. 4E**). Numerical simulations confirm that the cross-homeostatic rule robustly guides Up-states to different *E*_*set*_ and *I*_*set*_ setpoints (**Fig. 4F**), whose ratios define the slopes of the final relationship between the weights (**Fig. 4G**).

To further validate the effectiveness and stability of the cross-homeostatic rule we again used analytic methods to determine the eigenvalues of the 4-dimensional learning-rule dynamical system governed by the family of four cross-homeostatic rules. As above, stability is determined by the sign of the real part of the eigenvalues of the system. It can be shown (see **Section 1.3** in the Supplementary Material) that this learning rule is stable for any set of parameter values, provided that the stability conditions of the neural subsystem are satisfied (**Fig. 4H**). Therefore, it is possible to formally establish that the cross-homeostatic learning rules are inherently stable, and can robustly account for the emergence and maintenance of self-sustained inhibition-stabilized dynamics in the two-population model.

### Cross-homeostatic rules drive average activity in a multi-unit model to setpoints

The previous results demonstrate the robustness of the cross-homeostatic family of rules in driving a two-subpopulation rate model to a stable Up-state. We next examined if these rules are also effective when considering a multi-unit model in which there are many excitatory and inhibitory units. The firing-rate model was composed of 80 excitatory and 20 inhibitory recurrently connected neurons (**Fig. 5A**). In this case, individual neurons adjust their weights to minimize the average error of their presynaptic partners (see Methods). Starting with a random weight initialization, the network reaches stable self-sustained dynamics (**Fig. 5B-C**). However, individual units converge to different final rate values, satisfying the defined setpoints only as an average (green and red thick lines of **Fig. 5B**). This is a result of the nature of the cross-homeostatic rules: neurons adjust their weights to minimize the error of the *mean* activity of its presynaptic partners. For this reason, although the network is globally balanced, single units do not converge to the same balanced E-I line attractor (**Fig. 5D-E**), and little structure is observed in the connectivity matrix after learning (**Fig. 5F**). Simulations across 400 different initialization conditions demonstrate that the rules lead the average excitatory and inhibitory population activity to *E*_*set*_ and *I*_*set*_, respectively (**Fig. 5G-H)**. The cross-homeostatic rules are thus capable of driving a multi-unit model to a stable Up-regime, but they do not guide individual units to local setpoints.

**Figure 5.**
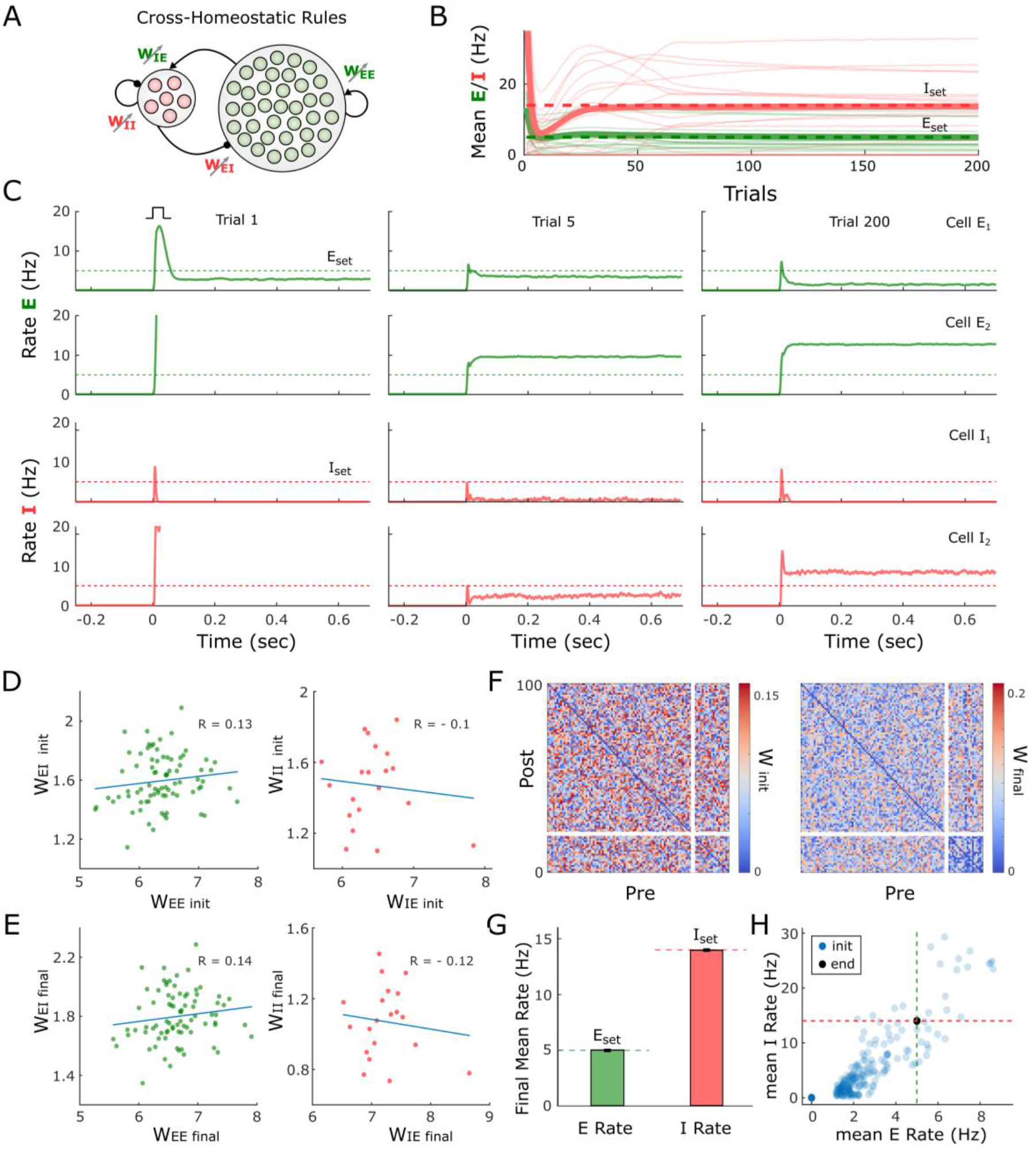
Cross-homeostatic rules drive a multi-unit firing rate model to a global network balance. **(A)** Schematic (left) of the multi-unit rate model. The network is composed of 80 excitatory and 20 inhibitory units recurrently connected. The four weight classes are governed by cross-homeostatic learning rules (right). See Methods for a detailed explanation of the implementation. **(B)** Evolution of the average rate across trials of 20 excitatory and inhibitory units in an example simulation. The network is initialized with random weights (see Methods) and so neurons present diverse initial rates. *E*_*set*_=5 and *I*_*set*_=14 represent the target homeostatic setpoints. Red and green lines represent the individual (thin lines) and average (thick lines) firing rate of inhibitory and excitatory population, respectively. **(C)** Example of the firing rate of two excitatory and two inhibitory units at different points in (B). The evolution of the firing rate of the excitatory and inhibitory population within a trial in response to a brief external input is shown in every plot. Individual units converge to a stable Up-state but not to the defined setpoint. **(D)** E-I weight relationships at the beginning of the simulation. Every dot represents the total presynaptic weight onto a single unit. Left excitatory neurons. Right inhibitory neurons. **(E)** Same plot as in D at the end of the simulation. **(F)** Weight matrix for the multi-unit model at the beginning (left) and end (right) of the simulation. First 20 neurons are inhibitory. **(G)** Average firing rate of the units of the multi-unit model and for different initializations of weights (n=400). The network converges to the setpoints in average. Data represents mean ± SEM. **(H)** Same data as in (G) but showing the average initial rate of the network for the multiple initializations (blue dots) and the average rate at the end (black). Target rates are shown in dotted lines (green, *E*_*set*_=5, red *I*_*set*_=14).

### A learning rule with cross-homeostatic and homeostatic terms leads to local convergence to setpoints

The above results demonstrate a potential limitation of the cross-homeostatic family of rules: the target setpoints are only reached at the population level. An additional and potentially more serious limitation is that cross-homeostatic rules predict that artificially altering the activity of a small number of excitatory neurons within a large network would not directly produce homeostatic plasticity in these neurons, but directly produce plasticity in their postsynaptic inhibitory neurons. This prediction seems to conflict with homeostatic plasticity experiments that have targeted specific cell types rather than globally alter activity through pharmacological means (Burrone et al., 2002; Xue et al., 2014). We therefore assessed the scenario in which both cross-homeostatic and homeostatic rules operate in parallel, resulting in a “two-term cross-homeostatic” family of rules. These rules can actually be recovered after an approximation of a gradient descent derivation on a loss function that includes the difference between *E* and *I* and their respective setpoints (see **Section 3** in the Supplementary Material). In a two-subpopulation model, we first confirmed that this two-term cross-homeostatic family is stable—assuming that the learning rate of the homeostatic term does not dominate (see Supplementary Material, **Section 1.4**). Simulations with the same multi-unit model as **Fig. 5** show that with the two-term cross-homeostatic rule all individual units converge to their respective *E*_*set*_ and *I*_*set*_ (**Fig. 6A-C**). Importantly, in contrast to the single-term cross-homeostatic rule the total excitatory and inhibitory weight of each individual unit converged to the E-I balance of the line attractor predicted by the network equations (**Fig. 6D-E**), while more structure is also observed in the weight matrices (**Fig. 6F**)—i.e., there is less homogeneity between the four synapse classes. The convergence to the setpoints was stable across a wide range of initial states (Fig. **6G-H**). Thus, a hybrid family of learning rules that includes both cross-homeostatic and homeostatic forces provide global network stability, while also locally driving each unit to their setpoint and a balanced E-I regime.

**Figure 6.**
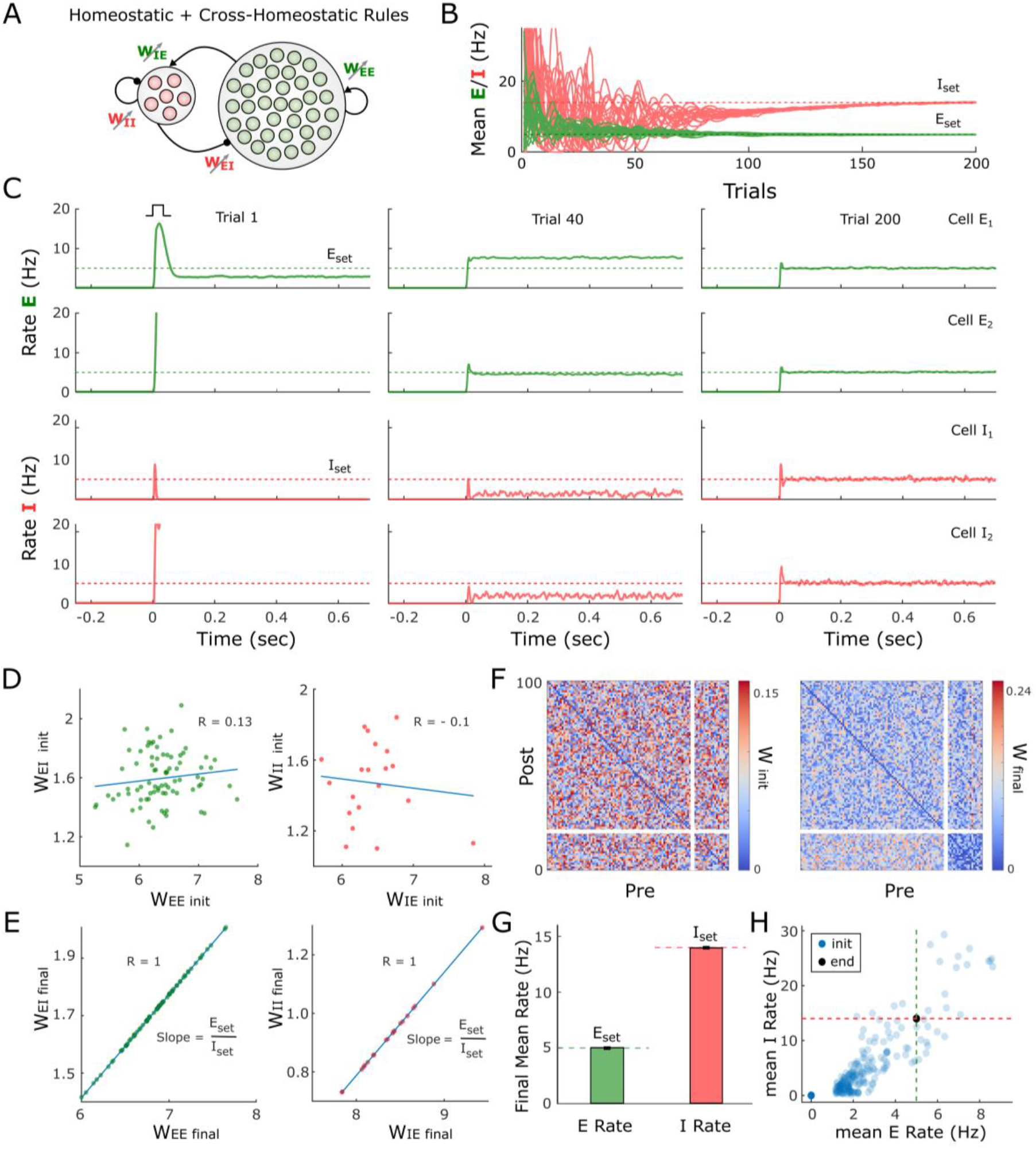
Adding cross-homeostatic influences to homeostatic rules lead global and local convergence to setpoints. **(A)** Schematic (left) of the multi-unit rate model. The network is composed of 80 excitatory and 20 inhibitory units recurrently connected. The four weight classes are governed by homeostatic rules with cross-homeostatic influences (right). See Methods for a detailed explanation of the implementation. **(B)** Evolution of the average rate across trials in an example simulation (20 excitatory and inhibitory units). The network is initialized with random weights (same as in **Fig. 5**, see Methods) and so neurons present diverse initial rates. *E*_*set*_=5 and *I*_*set*_=14 Hz represent the target homeostatic setpoints. **(C)** Example of the firing rate of two excitatory and two inhibitory units at different points in (B). The evolution of the firing rate of the excitatory and inhibitory population within a trial in response to a brief external input is shown in every plot. Units converge to a stable Up-state and at an individual setpoint. **(D)** E-I weight relationships at the beginning of the simulation. Every dot represents the total presynaptic weight onto a single unit. Left excitatory neurons. Right inhibitory neurons. **(E)** Same plot as in D at the end of the simulation. The network has reached a stable state and weights converge to single E-I balance defined by a line attractor. **(F)** Weight matrix for the multi-unit model at the beginning (left) and end (right) of the simulation. First 20 neurons are inhibitory. **(G)** Average firing rate of the units of the multi-unit model and for different initializations of weights (n=400). Data represents mean ± SEM. **(H)** Same data as in (G) but showing the average initial rate of the network for the multiple initializations (blue dots) and the average rate at the end (overlapping black circles). Target rates are shown in dotted lines (green, *E*_*set*_; red, *I*_*set*_).

## DISCUSSION

Elucidating the learning rules that govern the connectivity within neural circuits represents a fundamental goal in neuroscience, in part, because learning rules establish unifying principles that span molecular, cellular, systems, and computational levels of analyses. Elucidation of Hebbian associative synaptic plasticity, for example, linked simple computations at the level of single proteins (the NMDA receptor) with higher-order computations at the systems and computational levels (Hebb, 1949; Miller et al., 1989; Buonomano and Merzenich, 1998; Martin et al., 2000; Song et al., 2000). However, it remains the case that relatively little is known about the learning rules that give rise to complex neural dynamic regimes. Here we have taken steps towards exploring families of learning rules that operate in parallel at four different synapse classes and capture the experimentally observed emergence of Up-states in cortical networks.

Towards this goal we first confirmed that, in agreement with previous studies (Johnson and Buonomano, 2007; Motanis and Buonomano, 2020), Up-states emerge over the course of weeks of *ex-vivo* development. Because cortical organotypic cultures maintain much of their local and laminar architecture, and are mostly isolated from other cortical and subcortical inter-areal connectivity (Bolz, 1994; Echevarria and Albus, 2000; De Simoni et al., 2003), these results suggest the presence of local learning rules that lead to self-sustained, inhibition-stabilized activity in the absence of any supervisory, modulatory, or structured input signals from other brain areas.

We first explored whether standard formulations of homeostatic plasticity can account for the unsupervised emergence of Up-states—or more generally of self-sustained, inhibition-stabilized regimes. Based on experimental data we assumed that both excitatory and inhibitory neurons have an ontogenetically programmed activity setpoint during Up-states and that plasticity at the four weight classes is driven by homeostatic plasticity. Numerical simulations and analytical stability analyses revealed that while some initial conditions and parameter regimes led to self-sustained dynamics, they occupied a relatively narrow region of parameter space: when the rate of synaptic plasticity onto inhibitory neurons is much lower than that onto excitatory neurons (**Fig. 2**, and Supplementary Materials). When the rate of inhibitory and excitatory plasticity are comparable, analytical stability analyses confirmed that the region of stability of the network dynamics only overlapped in a narrow region. Such a narrow stability area seems incompatible with the robustness necessary in biological systems, and with experimental data showing that inhibitory neurons exhibit as much or more homeostatic plasticity than excitatory neurons (Keck et al., 2011; Kuhlman et al., 2013; Gainey et al., 2018; Ma et al., 2019). We thus conclude that a family of standard homeostatic learning rules operating at all four synapse classes is not sufficient to account for the experimentally observed emergence of self-sustained dynamics in cortical circuits.

### Cross-homeostatic plasticity

Analyses of approximations of a gradient-descent-derived learning rule suggested, somewhat counterintuitively, that adjusting the *E* population based on the error of the *I* population (and vice-versa) may prove to be an effective family of learning rules. Indeed, numerical simulations and analytical stability analyses revealed that this cross-homeostatic rule was robustly stable (**Fig. 4**). The convergence to the excitatory and inhibitory setpoints, however, only occurred at the population level, not at the level of individual units. This observation, however, is not inconsistent with experimental data, which shows that *in vivo* neurons do exhibit a wide range of variability in their apparent setpoints (Hengen et al., 2016; Trojanowski et al., 2020). However, a significant concern with this single-term cross-homeostatic rule is that it predicts that selectively increasing activity in a subpopulation of excitatory neurons would first induce plasticity in inhibitory neurons (*W*_*I*←*E*_ and *W*_*I*←*I*_)—which could in turn lead to plasticity in the manipulated excitatory neurons (*W*_*E*←*E*_ and *W*_*E*←*I*_). Most homeostatic plasticity studies do not speak to this prediction because they have used pharmacological manipulations of both excitatory and inhibitory neurons. However, some studies have used cell-specific manipulations—e.g., cell-specific overexpression of potassium channels (Burrone et al., 2002; Xue et al., 2014)—that strongly support the notion that synaptic plasticity is guided at least in part on their own deviation from setpoint.

In our opinion, and although we have explored alternative rules (see Supplementary Material, **Section 1.6**), the most biologically plausible set of learning rules that lead to stable Up-states comprises a hybrid rule that includes both standard homeostatic and cross-homeostatic terms. Such a two-term cross-homeostatic rule robustly led to a self-sustained, inhibition-stabilized network, led to all units converging to their setpoints, and is directly consistent with current experimental data.

### Biological plausibility of cross-homeostatic plasticity

While the neural mechanisms underlying homeostatic plasticity remain to be elucidated, it is generally assumed that an individual neuron can maintain a running average of their firing rate over the course of hours as a result of Ca^2+^-activated sensors. Based on the deviation of this value from an ontogenetically determined setpoint, neurons up- or down-regulate the density of postsynaptic receptors accordingly (Liu et al., 1998; Joseph and Turrigiano, 2017; Trojanowski et al., 2020). Two-term cross-homeostatic plasticity would require an additional, and apparently non-local information about the error in a given neuron’s presynaptic partners. It is important to stress, however, that this rule is not necessarily a non-local rule, because any postsynaptic neuron has access to the mean activity of its presynaptic partners simply as a result of its postsynaptic receptor activation. Indeed, a plasticity rule for *W*_*I*←*E*_ weights with a similar cross-homeostatic error term has also been recently proposed and implemented based on the mean activation of postsynaptic receptors—more specifically the net postsynaptic currents which provide a coupled measure of average presynaptic firing and synaptic weights (Mackwood et al., 2021).

Here we propose that cross-homeostatic plasticity could be implemented through metabotropic postsynaptic metabotropic receptors—e.g., mGlu and GABA_b_. Such receptors would provide a mechanism for postsynaptic neurons to maintain a running average of the activity of its presynaptic partners that is decoupled from the synaptic weights. Metabotropic receptors are G-protein coupled receptors (GPRC) that provide a low-pass filtered measure of presynaptic activity and are involved in a large number of incompletely understood neuromodulatory roles (Blein et al., 2000; Niswender and Conn, 2010). Since metabotropic receptors appear to undergo less homeostatic and associative plasticity, they provide a measure of presynaptic activity that is naturally decoupled from the ionotropic receptors (e.g., AMPA and GABA_a_) that are being up- and down-regulated.

Further support for the notion that individual neurons have access to global network activity emerges from studies suggesting that neurons might not homeostatically regulate activity at the individual neuron level, but rather at the global population level (Slomowitz et al., 2015). Such a global-level homeostasis could be achieved by non-synaptic paracrine transmission. Indeed, retrograde messenger systems are ideally suited for this role, as they have already been implicated in signaling mean activity levels to local capillaries, driving the activity-dependent vasodilation that underlies fMRI (Drew, 2019).

### The paradoxical effect and standard homeostatic rules

The paradoxical effect is one of the defining features of inhibition-stabilized networks, and a growing body of evidence suggests that Up-states and other self-sustained dynamic regimes are instantiations of inhibition-stabilized networks (Zucca et al., 2017; Mahrach et al., 2020; Sanzeni et al., 2020; Sadeh and Clopath, 2021). Here we show that the paradoxical effect applies important constraints to the potential learning rules that lead to the emergence of inhibition-stabilized networks. In the simplified case in which there is only homeostatic plasticity onto the inhibitory neurons, we can immediately see why the paradoxical effect renders standard homeostatic rules ineffective. If the *I* population is below its setpoint, the standard homeostatic rules would increase *W*_*I*←*E*_, which paradoxically would further decrease *I* (**Fig. 3**), thus further increasing the error instead of decreasing it (**Fig. 3**). This reasoning is related to why, when using the standard family of homeostatic rules, the rate of plasticity onto the inhibitory neurons has to be much smaller—in effect dampening the “paradoxical homeostatic plasticity effect”. Furthermore, our analytical stability analyses show that in the limit of vanishingly small excitatory learning rates (α_EE,EI_ << α_IE,II_) the stability region of the weight subsystem is bounded by the paradoxical condition. This means that the only allowed stable states with non-zero *E* activity will occur in the non-paradoxical regime, if any, and they will not be proper inhibition-stabilized Up-states.

### Future directions and experimental predictions

While we implemented homeostatic learning rules at all four synapses classes in our model, it is important to stress that we have omitted other well-characterized forms of synaptic plasticity. Of particular relevance, we did not include associative LTP or STDP. These forms of plasticity are generally considered to capture the correlation structure in networks which are driven by structured inputs. Arguably, because our circuits develop in the absence of any structured external input and because all excitatory and inhibitory neurons synchronously shift between Down↔Up states, it is possible that associative forms of plasticity do not contribute significantly to Up-state development. Nevertheless, future experimental and theoretical studies have to address the potential role for associative forms of synaptic plasticity in Up-state development.

An important implication of our results is that neuronal and network properties can operate in fundamentally different ways. That is, while homeostatic plasticity can lead to single neurons to reach their target setpoints in simple feedforward circuits, those same rules can be highly unstable when the neurons are placed even in the simplest of recurrent excitatory/inhibitory circuits with emergent dynamics. Furthermore, because emergent neural dynamic regimes are highly nonlinear, and in particular, that stable self-sustained dynamic regimes exhibit a paradoxical effect, it is likely that the brain exhibits “paradoxical” or counterintuitive learning rules to generate self-sustained dynamic regimes.

## METHODS

### Ex vivo slice preparation

Organotypic slices were prepared using the interface method (Stoppini et al., 1991; Goel and Buonomano, 2016). Briefly, five to seven day-old WT and PV-Cre mice were anesthetized with isoflurane and decapitated. The brain was removed and placed in chilled cutting media. Coronal slices (400 µm thickness) containing auditory cortex were sliced using a vibratome (Leica VT1200) and placed on filters (MillicellCM, Millipore, Billerica, MA, USA) with 1 mL of culture media. Culture media was changed at 1 and 24 hours after cutting and every 2-3 days thereafter. Cutting media consisted of EMEM (MediaTech cat. #15-010) plus (final concentration in mM): MgCl_2_, 3; glucose, 10; HEPES, 25; and Tris-base, 10. Culture media consisted of EMEM plus (final concentration in mM): glutamine, 1; CaCl_2_, 2.6; MgSO_4_, 2.6; glucose, 30; HEPES, 30; ascorbic acid, 0.5; 20% horse serum, 10 units/L penicillin, and 10 μg/L streptomycin. Slices were incubated in 5% CO2 at 35°C.

### Two-photon Calcium Imaging

Organotypic slices from PV-Cre mice (Jackson Laboratory #017320) were transfected at 1-3 DIV with pENN-AAV-CaMKII-GCaMP6f-WPRE-SV40 to selectively express GCaMP6f in excitatory neurons, and pAAV-CAG-Flex-mRuby2-GSG-P2A-GCaMP6f-WPRE-pA to visualize and selectively express GCaMP6f in PV^+^ inhibitory neurons. Transfection was achieved by gently delivering 1µL of each virus onto the slice using a micropipette. Experiments were performed at least 12 days after transfection to allow for robust expression.

Calcium imaging was performed with a galvo-resonant-scanning two-photon microscope (Neurolabware) controlled by Scanbox acquisition software (https://scanbox.org). A Coherent Chameleon Ultra II Ti:sapphire laser (Cambridge Technologies) was used for GCaMP6f (920 nm) and mRuby excitation (1040 nm). A 16x water-immersion lens (Nikon, 0.8 NA, 3 mm working distance) was used. Image sequences were captured using unidirectional scanning at a frame rate of ∼ 15 Hz. The size of the recorded imaging field was ∼ 520 × 800 µm (512 × 796 pixels). Five min of spontaneous activity was recorded at 920 nm at every developmental time point. Before the recording a snapshot at 1040 nm was recorded in order to identify PV^+^ neurons. Regions of interests (ROI) for both excitatory and PV^+^ neurons were established using the imaging processing pipeline Suite2p (https://github.com/MouseLand/suite2p) (Pachitariu et al., 2017). ΔF/F was calculated as (F(t) − F0))/F0, where F(t) was the raw fluorescence filtered with a median filter with a window of 1 s. F0 was the running median F(t) over the previous 20 s window. For each recorded slice and neural population (excitatory or PV^+^), potential Up-states were identified based on a threshold set at 1 above the mean z-scored raw fluorencence F trace of all neurons. If these events remained above threshold for at least one second, they were classified as Up-states. Mean Up Frequency and Duration were computed over all Up-states detected within the 5 min spontaneous activity period.

### Computational model

A two-population firing-rate model was implemented based on Jercog et al (2017). The firing rate of the excitatory (*E*) and inhibitory (*I*) population obeyed Wilson and Cowan dynamics (Wilson and Cowan, 1972):

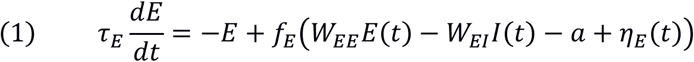

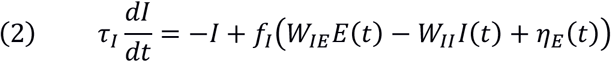

where *W*_*XY*_ represents the weight between the presynaptic unit *Y* and postsynaptic unit *X. τ*_*X*_ and *η*_*X*_ represent a time constant and an independent noise term, respectively. The time constants were set to *τ*_*E*_ = 10*ms* for the excitatory and *τ*_*I*_ = 2*ms* for the inhibitory subpopulations. The noise term was an Ornstein-Uhlenbeck process with mean *μ*_*x*_ = 0, a time constant 1/*θ*_*x*_ = 1*ms*, and a sigma parameter of *σ*_*x*_ = 10. To elicit Up-states a step current was injected at the beginning of each trial on the excitatory population.

The function *f*_*Y*_(*x*) represents the intrinsic excitability of the neurons, and it is modeled as a threshold-linear (ReLU) function with threshold *θ*_*Y*_ and gain *g*_*Y*_.

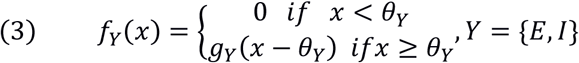

As in Jercog et al (2017) the thresholds were set to *θ*_*E*_ = 4.8 and *θ*_*I*_ = 25, and the gains to *g*_*E*_ = 1 and *g*_*I*_ = 4. The higher thresholds in PV neurons are consistent with experimental findings (Romero-Sosa et al., 2021).

The linear relationship between excitatory and inhibitory weights (**Fig. 4**) correspond to the steady-state solution of the neural subsystem when the inhibitory and excitatory rates are at its target setpoints. The solution can be obtained by setting the left side of equations (1) and (2) to zero, and substituting the steady state *E* and *I* values by *E*_*Set*_ and *I*_*Set*_.

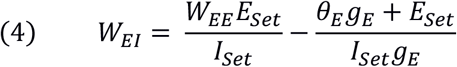

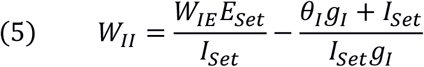

Thus, the slope of the E/I balance line in **Fig. 4** corresponds to *E*_*Set*_/*I*_*Set*_. See details and analytical results in **Section 2.2** of the Supplementary Material.

### Synaptic plasticity

Plasticity at all four weight classes (*W*_*E*←*E*_, *W*_*E*←*I*_, *W*_*I*←*E*_, *W*_*I*←*I*_) was governed by different families of homeostatic based learning rules, all driven by the deviation of the actual excitatory and inhibitory rates from their target setpoints (*E*_*set*_ and *I*_*set*_). Three different learning rules are presented in the main text of this paper.

#### Standard homeostatic family of rules

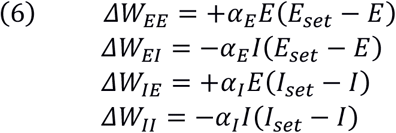

where *α*_*E*_ and *α*_*I*_ are the learning rates onto the excitatory and inhibitory units, respectively. All alphas are set to equal values in the simulation data shown in **Fig. 2** (*α* = 0.0001). The setpoints were based on empirically measured values in *ex vivo* cortical circuits (Romero-Sosa et al., 2021): *E*_*set*_ = 5 and *I*_*set*_ = 14 Hz.

The configuration of setpoints follows a classic homeostatic formulation (Turrigiano et al., 1998; Rossum et al., 2000; Liu and Buonomano, 2009; Vogels et al., 2011), where every neural population adapts its input weights homeostatically in order to minimize its error term. As outlined in Supplementary Material (**Section 1.5**) we also examined variants of this formulation, such as standard synaptic scaling (which includes the weight as factor).

We prove that these rules are only stable in a narrow parameter regime (when excitatory plasticity dominates). See details and analytical results in **Section 2** of the Supplementary Material.

#### Single-term cross-homeostatic family of rules

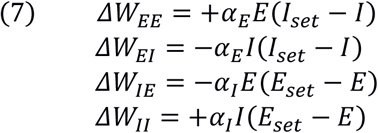

All alphas are set to equal values in the simulation data shown in **Fig. 4** (*α* = 0.0001), except on the example shown in **Fig. 4B-C**, where a rate of *α* = 0.0005 was used. We note that an equivalent rule for *W*_*IE*_ has been recently derived (Mackwood et al., 2020). For **Fig. 4** two alternative pairs of setpoints were explored (*E*_*set*_ = 5 and *I*_*set*_ = 24) and (*E*_*set*_ = 10 and *I*_*set*_ = 14).

We prove that these rules are stable for any set of parameters. See details and analytical results in **Section 1.3** of the Supplementary Material.

#### Two-term cross-homeostatic family of rules

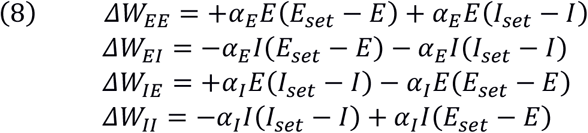

A single shared learning rate (*α* = 0.00001) was used for **Fig. 6** (see section below on the multi-unit model). We prove that these rules are stable for a biologically meaningful set of parameter values, as long as the homeostatic part does not dominate (Supplementary Material, **Section 1.4**). The two-term rules combine homeostatic and cross-homeostatic terms. This formulation can be obtained after an approximation of a gradient descent derivation on the following loss function:

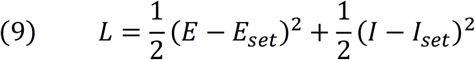

The mathematical derivation can be found in the Supplementary Material (**Section 3**).

An additional *Forced-Balance* learning rule, which exploits the steady-state solution of the neural subsystem at its target setpoints (equations 5 and 6), has also been explored (see **Section 1.6** of the Supplementary Material).

#### All rules, numerical simulations

For all simulations, the weights were updated after the completion of every trial. The trials lasted 2 seconds. For our numerical simulations, *E* and *I* on every rule are implemented as average firing rates. The average of *E* and *I* is computed after every trial and then is low pass filtered by a process with a time constant *τ*_*trial*_ = 2. The numerical integration time step was 0.1 ms. A minimum weight of 0.1 was set for all weights.

A saturation to the excitatory and inhibitory firing rate (100 and 250 Hz, respectively) was added to prevent the nonbiological scenario in which activity could diverge towards infinity under unstable conditions. Note the saturation is not necessary for the cross-homeostatic rule because it is inherently stable as proved in the Supplementary Material (**Section 1.3**).

In **Fig. 2D** and **4D-G** we initialize the weights uniformly in between the following ranges: *W*_*EE*_[4,7], *W*_*EI*_[0.5,2], *W*_*IE*_ [7,13], *W*_*II*_[0.5,2]. Simulations were run for 3000 trials to assess stability and convergence.

#### All rules, analytical stability analyses

We analyzed the entire dynamical system (composed of the neural subsystem and learning rule subsystem) for every synaptic learning rule considered in this work, and analyzed its stability. In every case, the general prescription is:

a. Take the combined neural and learning rule subsystems and nondimensionalize all variables, so that the two different time scales are evident (fast neural, slow synaptic plasticity). For the description of the learning rule subsystem we switch from discrete-time dynamics to continuous-time dynamics: ΔW → τ_0_ dW/dt
b. Make a quasi-steady state (QSS) approximation of the neural subsystem. This means we will consider the neural subsystem is fast enough so that it converges “instantaneously” (when compared to the synaptic plasticity subsystem) to its corresponding fixed point. For this we will require that the stability conditions of the neural subsystem are satisfied (see below).
c. Find the steady-state solution of the synaptic plasticity subsystem, i.e. the Up-state fixed point; compute the Jacobian of the synaptic plasticity subsystem at the Up-state; compute the eigenvalues of the Jacobian. Two out of the four eigenvalues are expected to be zero because the Up-state is not an isolated fixed point of the system but a continuous 2D plane in 4D weight space.
d. Address (linear) stability. If both nonzero eigenvalues have negative real parts, then the Up-state is stable under the learning rule; if at least one of the nonzero eigenvalues has positive real part, then the Up-state is unstable. (A note on abuse of notation: we might say indistinctly “the Up-state is stable/unstable” and “the learning rule is stable/unstable”.)

See **Section 2** in the Supplementary Material.

### Multi-unit firing rate model

A rate-based recurrent network model containing *N*_*e*_ = 80 excitatory and *N*_*i*_ = 20 inhibitory neurons was implemented with all-to-all connectivity (without self-connections). The activation of the neurons followed equations (1), (2) and (4). The same parameters as for the population model were used, where *W*_*XY*_ represents now a matrix of synaptic weights from population *X* to population *Y*. A minimum weight of 0.1/*N*_*x*_ for *W*_*EI*_ and *W*_*IE*_ and 0.1/(*N*_*x*_-1) for *W*_*EE*_ and *W*_*II*_ was set for all weights.

The synaptic plasticity rules were implemented as follows.

#### Cross-homeostatic family of rules

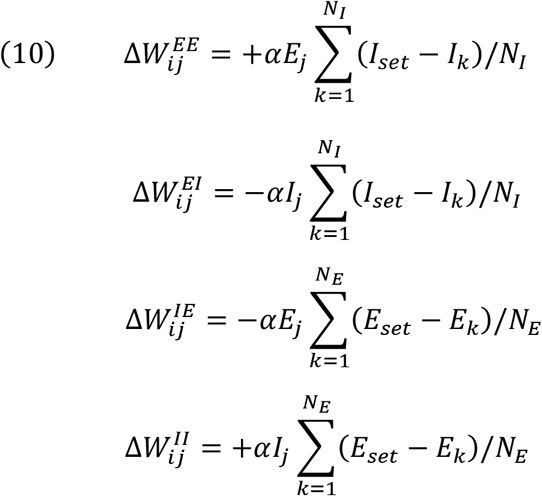

Where *i* and *j* represent the post- and presynaptic neurons, respectively, and *k* denotes the presynaptic inhibitory neurons targeting the excitatory neurons (or the presynaptic excitatory neurons targeting an inhibitory neuron). *N*_*E*_ and *N*_*I*_ denote the total number of excitatory and inhibitory neurons, respectively. The weights are therefore updated following the *average* presynaptic error of the crossed E/I population classes. Note as stated above that this formulation can be implemented in a local manner (see Discussion). A learning rate of *α* = 0.00002 was used for all simulations.

#### Two-term cross-homeostatic family of rules

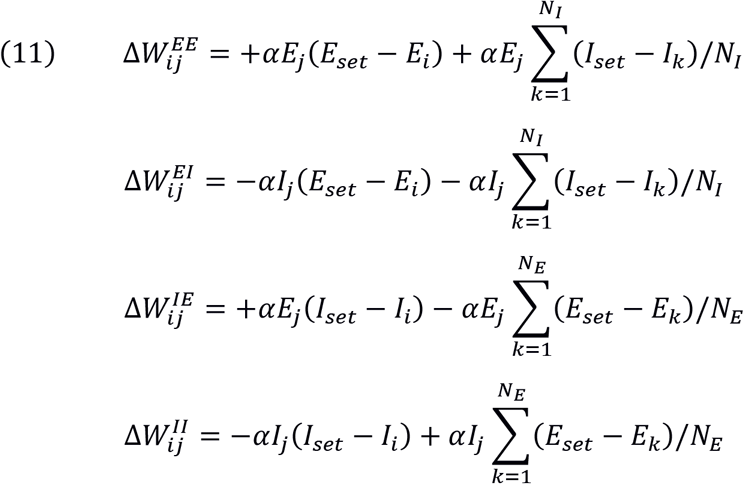

Here the first term represents the standard homeostatic rule, and the second term cross-homeostatic plasticity (as implemented above). A learning rate of *α* = 0.00001 was used for all simulations.

In **Fig. 5G-H** and **6G-H** we initialize the mean weights of the population uniformly in between the following ranges: *W*_*EE*_[1,6], *W*_*EI*_[0.5,2], *W*_*IE*_ [5,7], *W*_*II*_[0.5,2]. The weights within each class were normally distributed around that mean (normalized by the number of neurons) with a variance of 0.1. Note that this initialization led to multiple initial conditions with exploding network rates (which were held in check by the saturation cutoff). Those initial rates are not displayed in **Fig. 5-6H** for visualization purposes, but the rules successfully brought all those cases to the corresponding setpoints. Simulations were run for 1000 trials to assess stability of the convergence. In the example shown in **Fig. 5A-F** and **6A-F** the weights were initialized uniformly in the interval [0 0.16] and the simulation was run for 200 trials.

### Statistics and Software availability

Data are represented by the mean ± SEM. In **Fig. 1** a two-way ANOVA was performed to assess interaction of time and group (cell-type) on the development of Up-states.

Experimental and computational analysis were performed in custom-written MATLAB R2020a software. SageMath was used for the analytical proofs (see Supplementary Material). The MATLAB source code that reproduces **Fig. 2, 4, 5** and **6** is available at https://github.com/saraysoldado/UpDev2021. The Jupyter notebooks with SageMath code to reproduce all analytical results are available at: https://github.com/SMDynamicsLab/UpDev2021.

## ACKNOWLEDGEMENTS

We thank Juan Romero-Sosa and Helen Motanis for the sample traces shown in Figure 1. We thank Ben Liu, Shanglin Zhou, and Helen Motanis, for technical assistance and helpful discussions. We thank Joana Soldado-Magraner and Mike Seay for comments on the manuscript. This research was supported by NIH grant NS116589, and SSM was supported by the Swiss National Science Foundation (P2ZHP3-187943).

## SUPPLEMENTARY FIGURES

**Figure S1.**
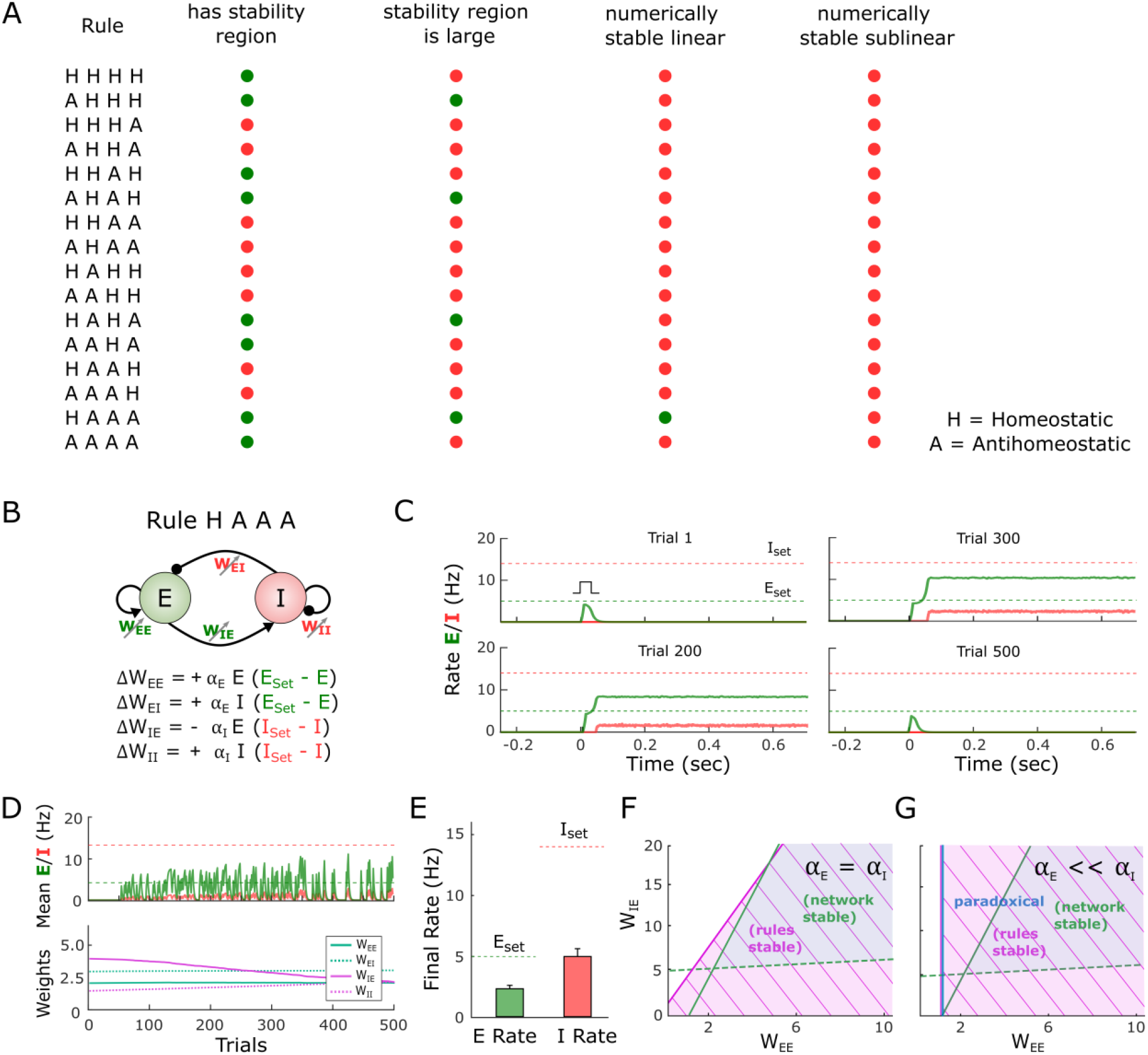
Homeostatic and anti-homeostatic combinations of learning rules also fail to generate the emergence of self-sustained dynamics. **(A)** Sixteen variations of the standard homeostatic rules presented in **Fig. 2** were assessed for stability. The learning governing each four weight types, *W*_*EE*_, *W*_*EI*_, *W*_*IE*_, *W*_*II*_ was set to be either homeostatic (H) or antihomeostatic (A). The first rule on the table (HHHH) corresponds to the standard homeostatic rules presented in **Fig. 2**, where all weights obey homeostatic learning. All rules were tested for stability analytically and numerically. A red dot implies that the listed condition is not satisfied, while a green dot means that it does. The condition on the first column indicates whether a stability region for the learning rule is present. The second column indicates whether such region has a large overlap with the region of stability of the neural subsystem. The third column indicates whether the rule is successful, using numerical simulations, at driving the network to a stable Up-state when starting from regimes with self-sustained activity already present (meaning the network is initialized in the linear regime). The fourth column indicates the same as the former, but with the network initialized in the sub-linear regime, where activity is not initially present (e.g., as observed early in developmental conditions). **(B)** Schematic (top) of the population rate model in which the four weights are governed by the HAAA rule in panel (A). **(C)** Example simulation of the HAAA rule over the course of simulated development. The evolution of the firing rate of the excitatory and inhibitory population within a trial in response to a brief external input is shown in every plot. *E*_*set*_*=5* and *I*_*set*_=14 represent the target homeostatic setpoints. Weights were initialized to *W*_*EE*_=2.1, *W*_*EI*_=3, *W*_*IE*_=4, and *W*_*II*_=2 as in **Fig. 2**. Note that while an evoked Up-state emerges by Trial 200 the firing rates do not converge to their setpoints, and by Trial 500 the Up-state is no longer observed. **(D)** Average rate across trials (upper plot) for the excitatory and inhibitory populations for the data shown in (C). Weight dynamics (bottom plot) produced by the homeostatic rules across trials for the data shown in (C). **(E)** Average final rate for 100 independent HAAA simulations with different weight initializations. Those initializations included cases in which the network starts in the sublinear regime (where the initial *E* firing rate was zero or very low). The weights were initialized uniformly between the following ranges: *W*_*EE*_ [1,3], *W*_*EI*_ [0.5,1.5], *W*_*IE*_ [4,8], *W*_*II*_[0.2,0.8]. Data represents mean ± SEM. **(F)** Analytical stability regions of the neural and HAAA learning rule subsystems as a function of the free weights *W*_*EE*_ and *W*_*IE*_. Here the stability plot is obtained by considering equal learning rates for all four learning rules (as used for panels C-E). **(G)** Similar to F but with but with *α*_*E*_ << *α*_*I*_. Right of blue line shows the area where the network is in a paradoxical regime (defined by the condition W_EE_ * g_E_ – 1 > 0). Contrary to standard homeostatic rules (**Fig. 2**), the HAAA rule is only stable in the paradoxical region of parameter space (i.e., W_EE*_g_E_ – 1 > 0; note white area to the left of the blue line). This may explain why the rule fails at driving the network to an Up-state when starting with developmental-like conditions.

