## SupplementaryMaterial for "Orchestrated Excitatory and Inhibitory Learning Rules Lead to the Unsupervised Emergence of Self-sustained and Inhibition-stabilized Dynamics"

September 10, 2021

### 1 Summary of results

In this section we describe the general approach and results of the analytical stability analyses of the joint neural and synaptic learning rules subsystems. We express results in terms of the "free weights" $W_{EE}$ and $W_{IE}$. In Section 2 we provide a detailed description of the approach. .

#### 1.1 ***Homeostatic*** learning rule

In continuous-time dynamics, the equations for the Homeostatic learning rule are

$$\begin{aligned}\frac{dW_{EE}}{dt} &= +\alpha_{EE}\,E(E_{set}-E)\\ \frac{dW_{EI}}{dt} &= -\alpha_{EI}\,I(E_{set}-E)\\ \frac{dW_{IE}}{dt} &= +\alpha_{IE}\,E(I_{set}-I)\\ \frac{dW_{II}}{dt} &= -\alpha_{II}\,I(I_{set}-I)\end{aligned} \tag{1}$$

The condition for the Up-state to be stable (i.e., the two nonzero eigenvalues to have negative real parts, see Section 2) under this rule is:

$$\begin{aligned}(E_{set}^2\alpha_{EE}+I_{set}^2\alpha_{EI})E_{set}g_E W_{IEup}&\\ >+(E_{set}^2\alpha_{IE}+I_{set}^2\alpha_{II})I_{set}(W_{EEup}g_E-1)&\\ +(E_{set}^2\alpha_{EE}+I_{set}^2\alpha_{EI})\Theta_I g_E&\end{aligned} \tag{2}$$

It is difficult to determine whether the stability condition of Eq. 2 is satisfied for a general set of parameter values (see numerical analysis below). However, this condition can be re-expressed in a more useful form in terms of $W_{EE}$ and $W_{II}$:

$$\begin{aligned}(R^2\alpha_3+\alpha_4)(W_{EEup}\,g_E-1)g_I&\\ <(R^2+\alpha_2)(W_{IIup}\,g_I+1)g_E&\end{aligned} \tag{3}$$

where

$$\begin{aligned}R&=E_{set}/I_{set}\\ \alpha_2&=\alpha_{EI}/\alpha_{EE}\\ \alpha_3&=\alpha_{IE}/\alpha_{EE}\\ \alpha_4&=\alpha_{II}/\alpha_{EE}\end{aligned}$$

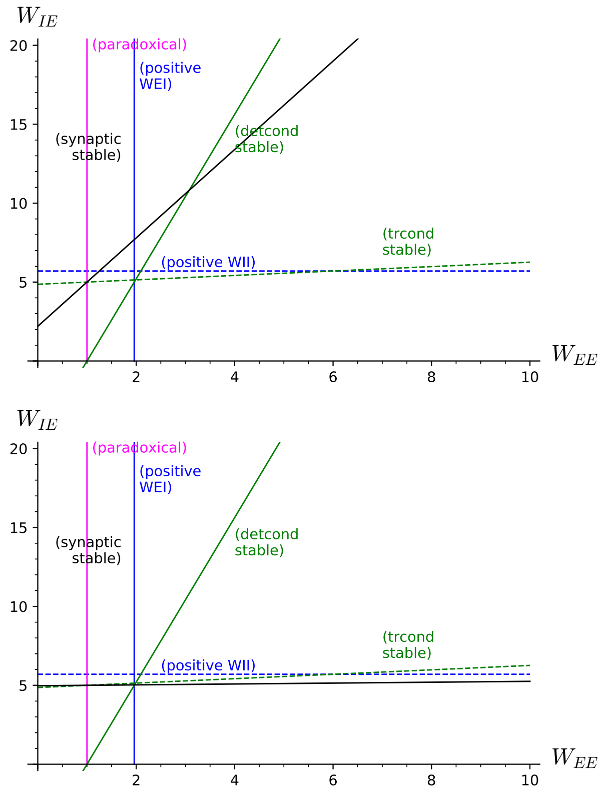

Figure S1: Regions of stability, *Homeostatic* learning rule. (Top) For biologically backed parameter values (Table 1) and learning rates of the same value ($\alpha_{XY} = 0.02$), the stability region of the Homeostatic learning rule has little overlap with the region where the neural subsystem is stable. (Bottom) Setting $\alpha_{EE} = \alpha_{EI} = 0.02$ and $\alpha_{IE} = \alpha_{II} = 0.0002$ enlarges the stability region of the learning rule and makes it overlap with the stability region of the neural subsystem. Every label is on the side where the corresponding condition holds (synaptic stable: Eq. 2; detcond stable: Eq. 22; trcond stable: Eq. 23; positive $W_{EI}$: Eq. 25; positive $W_{II}$: Eq. 26; paradoxical: Eq. 27).

Note that learning rate values of the same order lead to $\alpha_{2,3,4} \sim 1$ and that biologically backed parameter values satisfy:

$$I_{set} > E_{set}$$
$$g_I > g_E$$

both likely preventing the condition to hold. On the other hand, if $\alpha_{IE}$ and $\alpha_{II}$ are small enough (slow dynamics of the weights onto the inhibitory neuron) the rule can be stable. See the step-by-step derivation of this stability condition in Section 2.3.

As an illustration of the results above, in Figure S1(top) we plot the stability condition Eq. 2 with parameter values as in Table 1 and learning rates $\alpha_{XY} = 0.02$. It is clear that the learning rule is stable in a region with little overlap with the stability region of the neural subsystem. The stability region can be enlarged by making the dynamics of the weights onto the inhibitory neuron slower, as in Figure S1(bottom) where $\alpha_{EE} = \alpha_{EI} = 0.02$ and $\alpha_{IE} = \alpha_{II} = 0.0002$.

See Section 2.3 for a detailed analysis.

#### 1.2 Homeo-antiHomeo variations

The stability condition in the previous section was obtained by assuming all learning rates are positive. Interestingly, if some of them are negative then the Up-state may still be stable. A negative learning rate can be interpreted as the corresponding equation being *anti*-homeostatic, i.e. if the neural activity ($E$ or $I$) departs from its setpoint then the rule will drive it even farther away. While this kind of behavior would be usually deemed undesired, it is worth considering due to its relationship with the paradoxical regime.

In this section we consider the Homeostatic rule, Eq. 28, and let the learning rates $\alpha_{XY}$ be either positive or negative. The particular case where all learning rates are positive corresponds to the original Homeostatic learning rule.

Once we free the signs of the learning rates, the Up-state needs two conditions to be stable:

$$(R^2\alpha_3+\alpha_4)(W_{EE\,up}\,g_E - 1)g_I < (R^2+\alpha_2)(W_{II\,up}\,g_I + 1)g_E \tag{4}$$

$$(R^2\alpha_3+\alpha_4)(R^2+\alpha_2) > 0 \tag{5}$$

where

$$R = E_{set}/I_{set}$$
$$\alpha_2 = \alpha_{EI}/\alpha_{EE}$$
$$\alpha_3 = \alpha_{IE}/\alpha_{EE}$$
$$\alpha_4 = \alpha_{II}/\alpha_{EE}$$

Eq. 4 is equal to the stability condition of the original Homeostatic rule (Eq. 3). The additional condition Eq. 5 is very interesting in that it allows the Up-state to be stable, for instance, under a full anti-Homeo learning rule where all four learning rates are negative (leading to $\alpha_{2,3,4}$ all positive).

See details in the corresponding section of the SageMath-Jupyter notebook:
`upstates-Homeostatic stability.ipynb`

#### 1.3 *Cross-Homeostatic* learning rule

In continuous-time dynamics, the equations for the Cross-Homeostatic learning rule are

$$\begin{aligned} \frac{dW_{EE}}{dt} &= +\alpha_{EE}E(I_{set} - I) \\ \frac{dW_{EI}}{dt} &= -\alpha_{EI}I(I_{set} - I) \\ \frac{dW_{IE}}{dt} &= -\alpha_{IE}E(E_{set} - E) \\ \frac{dW_{II}}{dt} &= +\alpha_{II}I(E_{set} - E) \end{aligned} \tag{6}$$

and its stability condition in terms of the free weights $W_{EE}$ and $W_{IE}$ reads:

$$\begin{aligned} (E_{set}^2\alpha_{EE} + I_{set}^2\alpha_{EI})I_{set}g_E W_{IEup} \\ > -(E_{set}^2\alpha_{IE} + I_{set}^2\alpha_{II})E_{set}g_E W_{EEup} \\ + (E_{set}^2\alpha_{IE} + I_{set}^2\alpha_{II})(\Theta_E g_E + E_{set}) \end{aligned} \tag{7}$$

This stability condition can be put in a simpler form by switching to $W_{EI}$ and $W_{IE}$:

$$(R^2\alpha_3 + \alpha_4)W_{EIup} + (R^2 + \alpha_2)W_{IEup} > 0 \tag{8}$$

(where $R$ and $\alpha_{2,3,4}$ are defined as in the previous subsection). This condition is always satisfied because the weights and parameters are positive definite and thus the rule is stable for any choice of parameter values (as long as the neural subsystem is). Fig. S2 shows the stability region of the neural subsystem for the set of parameter values of Table 1. Any choice of values for the weights $W_{EE}$ and $W_{IE}$ within the stability region of the neural subsystem will lead to a stable Up-state.

See Section 2.4 for a detailed analysis.

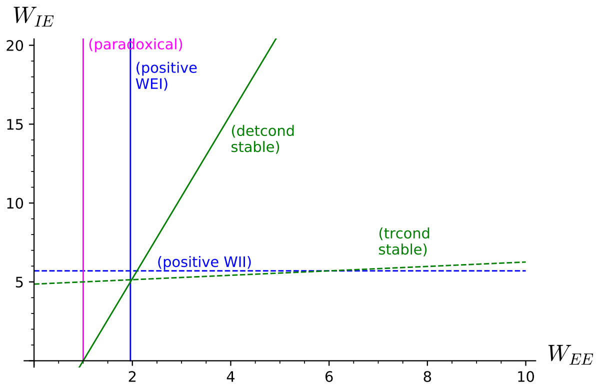

Figure S2: Stability of the *Cross-Homeostatic* rule. The rule is stable for any parameter value; the Up-state is thus stable where the neural subsystem is stable, i.e. in the upper right region between the two green lines. Every label is on the side where the corresponding condition holds (detcond stable: Eq. 22; trcond stable: Eq. 23; positive $W_{EI}$: Eq. 25; positive $W_{II}$: Eq. 26; paradoxical: Eq. 27). Parameter values as in Table 1.

#### 1.4 *Two-Term* learning rule

The equations for the Two-Term learning rule in continuous-time dynamics are

$$\begin{aligned} \frac{dW_{EE}}{dt} &= +\alpha E(I_{set} - I) + \beta E(E_{set} - E) \\ \frac{dW_{EI}}{dt} &= -\alpha I(I_{set} - I) - \beta I(E_{set} - E) \\ \frac{dW_{IE}}{dt} &= -\alpha E(E_{set} - E) + \beta E(I_{set} - I) \\ \frac{dW_{II}}{dt} &= +\alpha I(E_{set} - E) - \beta I(I_{set} - I) \end{aligned} \tag{9}$$

and its stability condition in terms of the free weights

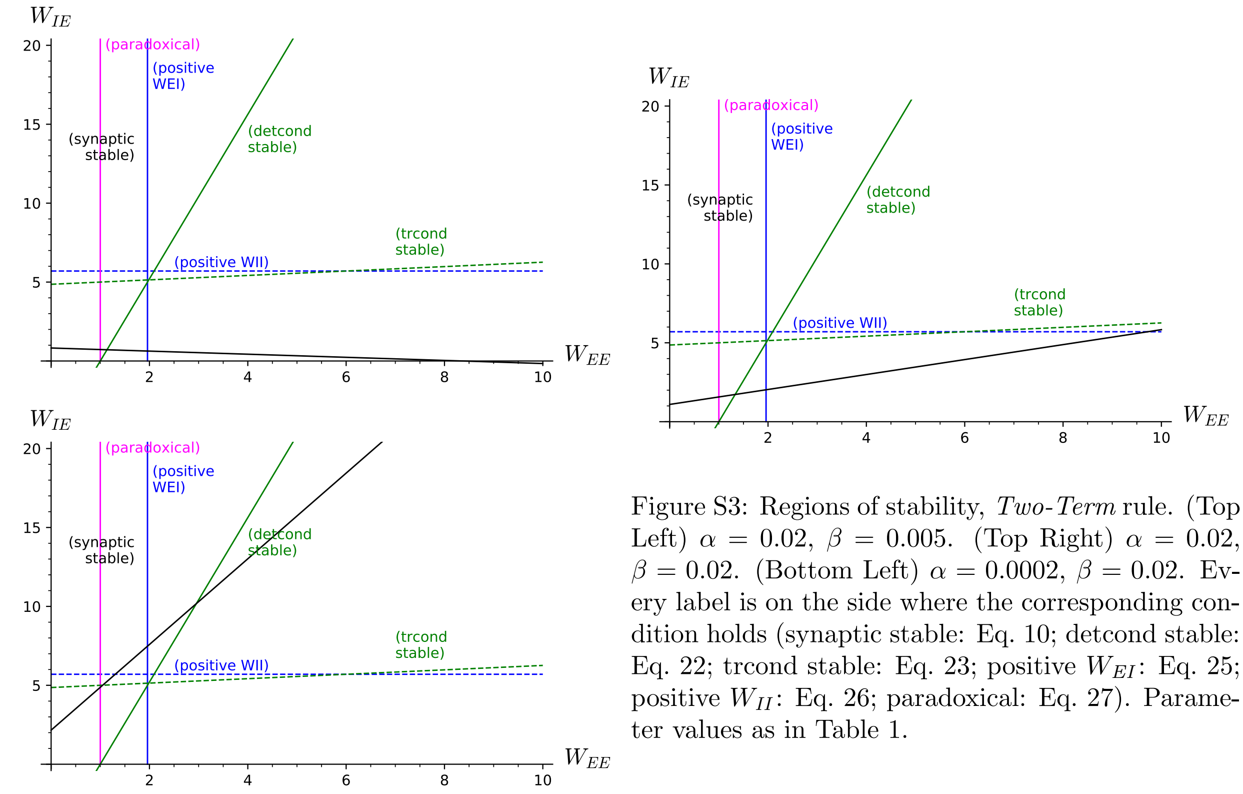

Figure S3: Regions of stability, *Two-Term* rule. (Top Left) $\alpha = 0.02$, $\beta = 0.005$. (Top Right) $\alpha = 0.02$, $\beta = 0.02$. (Bottom Left) $\alpha = 0.0002$, $\beta = 0.02$. Every label is on the side where the corresponding condition holds (synaptic stable: Eq. 10; detcond stable: Eq. 22; trcond stable: Eq. 23; positive $W_{EI}$: Eq. 25; positive $W_{II}$: Eq. 26; paradoxical: Eq. 27). Parameter values as in Table 1.

$W_{EE}$ and $W_{IE}$ is

$$\begin{aligned}(I_{set}\alpha + E_{set}\beta)g_E W_{IE\,up} \\ > (I_{set}\beta - E_{set}\alpha)g_E W_{EE\,up} \\ + (\Theta_E g_E + E_{set})\alpha + (\Theta_I g_E - I_{set})\beta\end{aligned} \tag{10}$$

In Figure S3 we plot the stability condition of this rule, Eq. 10, for three different parameter values: $\alpha \gg \beta$ (the "Cross-Homeostatic" terms dominate over the "Homeostatic" terms, and the rule is stable with the largest stability region); $\alpha = \beta$ (the two terms are of comparable size); and $\alpha \ll \beta$ (the "Homeostatic" terms dominate instead, and the stability region of the rule is as small as that of the Homeostatic learning rule).

In order to determine the validity of the stability condition, Eq. 10, in a more general situation, we rewrite it in a more useful form:

$$(a - b)\beta < (a' + b' + c)\alpha \tag{11}$$

where

$$\begin{aligned} a &= (W_{EE\,up} g_E - 1)E_{set} I_{set} g_I \\ a' &= (W_{EE\,up} g_E - 1)E_{set}^2 g_I \\ b &= (W_{II\,up} g_I + 1)E_{set} I_{set} g_E \\ b' &= (W_{II\,up} g_I + 1)I_{set}^2 g_E \\ c &= (I_{set}\Theta_I - E_{set}\Theta_E)g_E g_I \end{aligned}$$

Note that the following is satisfied for a biologically backed set of parameter values:

$$\begin{aligned} I_{set} &> E_{set} \\ \Theta_I &> \Theta_E \end{aligned}$$

and thus it is likely that $c > 0$. In addition, $b$ and $b'$ are positive definite, and $a, a' > 0$ in the paradoxical regime ($W_{EE} g_E - 1 > 0$). All this makes the stability condition likely satisfied, and thus the learning rule stable. Finally, a small enough $\beta$ would make the condition more likely to hold.

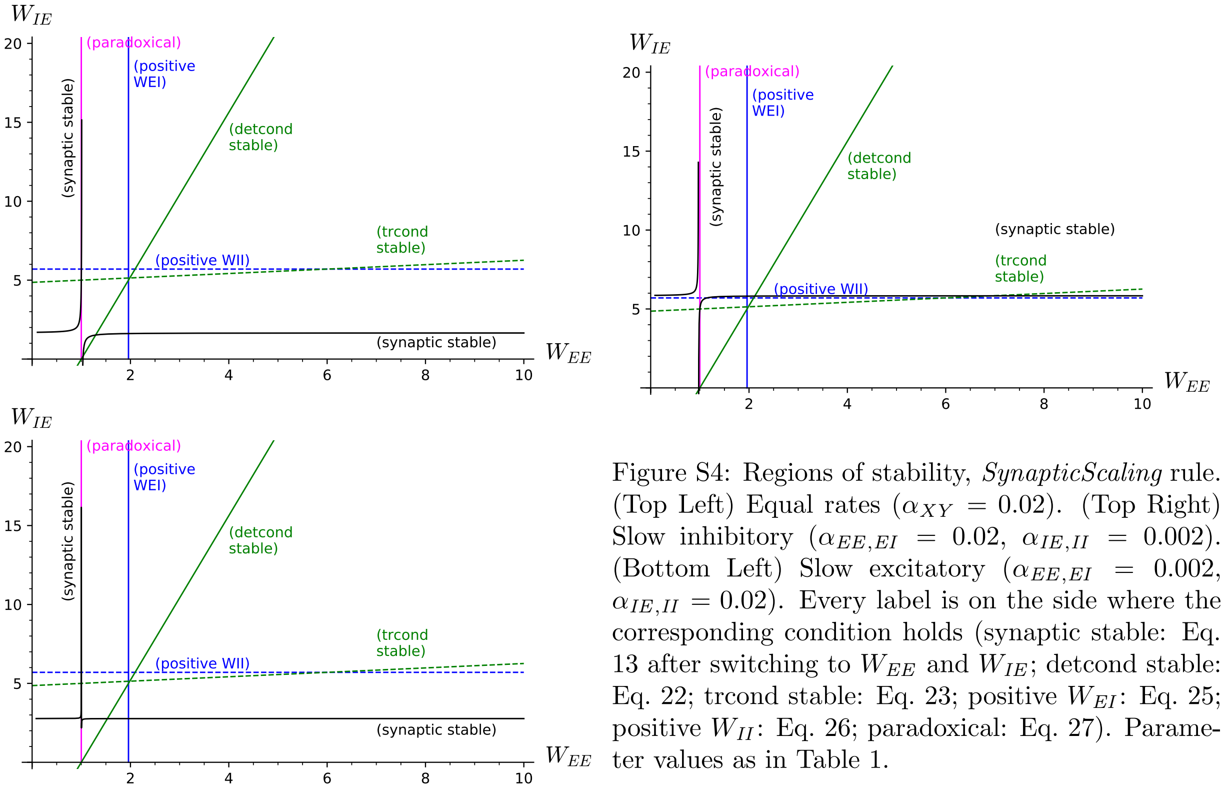

Figure S4: Regions of stability, *SynapticScaling* rule. (Top Left) Equal rates ($\alpha_{XY} = 0.02$). (Top Right) Slow inhibitory ($\alpha_{EE,EI} = 0.02$, $\alpha_{IE,II} = 0.002$). (Bottom Left) Slow excitatory ($\alpha_{EE,EI} = 0.002$, $\alpha_{IE,II} = 0.02$). Every label is on the side where the corresponding condition holds (synaptic stable: Eq. 13 after switching to $W_{EE}$ and $W_{IE}$; detcond stable: Eq. 22; trcond stable: Eq. 23; positive $W_{EI}$: Eq. 25; positive $W_{II}$: Eq. 26; paradoxical: Eq. 27). Parameter values as in Table 1.

See Section 2.4 for a detailed analysis.

#### 1.5 ***SynapticScaling*** learning rule

The equations for the SynapticScaling learning rule in continuous-time dynamics are

$$\begin{aligned} \frac{dW_{EE}}{dt} &= +\alpha_{EE}(E_{set} - E)W_{EE} \\ \frac{dW_{EI}}{dt} &= -\alpha_{EI}(E_{set} - E)W_{EI} \\ \frac{dW_{IE}}{dt} &= +\alpha_{IE}(I_{set} - I)W_{IE} \\ \frac{dW_{II}}{dt} &= -\alpha_{II}(I_{set} - I)W_{II} \end{aligned} \tag{12}$$

and the condition for the Up-state to be stable under this rule is

$$(W_{EE\,up}g_E - 1)a < (W_{II\,up}g_I + 1)b \tag{13}$$

where

$$\begin{aligned} a &= (I_{set}W_{II}\alpha_4 + \Theta_I\alpha_3)g_I \\ b &= E_{setup}W_{EEup}g_E \\ &\quad + ((W_{EEup}g_E - 1)E_{set} - \Theta_E g_E)\alpha_2 \\ &\quad - (W_{EEup}g_E - 1)I_{set}\alpha_3 \end{aligned}$$

(where $\alpha_{2,3,4}$ are defined as in previous subsections). This stability condition does not hold for biologically backed parameter values unless the dynamics of the weights onto the inhibitory neuron are slow enough (and in a few fine-tuned cases). To show this, we express the stability condition in terms of the free weights $W_{EE}$ and $W_{IE}$ and plot it with parameter values as in Table 1 and equal rates ($\alpha_{XY} = 0.02$; Figure S4 top left). The stability condition is a homographic function (i.e. a hyperbola) with stability regions in its upper-left and lower-right quadrants—entirely outside the stability region of the neural subsystem. If the dynamics of the weights onto the excitatory neuron are made slower, the homographic function is even steeper (bottom left); if the weights onto the inhibitory neuron are made slower instead, the stability regions switch and overlap with the stability region of the neural subsystem, making the Up-state stable (top left).

It is illustrative to consider the particular case where all learning rates are equal. In this case the stability condition, Eq. 13, doesn't depend on the learning rates and takes the simpler form:

$$\begin{aligned}(W_{IIup}g_I{+}1)a > (W_{EEup}g_E - 1)a' \\ + (W_{EEup}g_E - 1)(W_{IIup}g_I + 1)b\end{aligned} \tag{14}$$

where

$$\begin{aligned}a &= (E_{set}W_{EEup} - \Theta_E)g_E \\ a' &= (I_{set}W_{IIup} + \Theta_I)g_I \\ b &= I_{set} - E_{set}\end{aligned}$$

Note that in a biologically backed set of parameter values the following is true:

$$\begin{aligned}I_{set} &> E_{set} \\ g_I &> g_E \\ \Theta_I &> \Theta_E\end{aligned}$$

This makes $b > 0$ and likely $a' > a$ (in addition, $a'$ is a sum of positive terms while $a$ is a difference). Then in the paradoxical regime ($W_{EE}g_E - 1 > 0$) it seems likely that the stability condition is not satisfied, because the right-hand side is a sum of positive terms and one of them is likely greater than the left-hand side. The SynapticScaling rule is then likely unstable when the learning rates are equal.

A more general case with different learning rates can be analyzed by grouping terms in the following way:

$$\begin{aligned}(I_{set}W_{IIup}\alpha_4 + \Theta_I\alpha_3)g_I(W_{EEup}g_E - 1) \\ <(((W_{EEup}g_E - 1)E_{set} - \Theta_E g_E)\alpha_2 \\ - (W_{EEup}g_E - 1)I_{set}\alpha_3 \\ + E_{set}W_{EEup}g_E)(W_{IIup}g_I + 1)\end{aligned}$$

If $(W_{EE}g_E - 1) > 0$ (paradoxical regime), then decreasing $\alpha_3$ and/or $\alpha_4$ (slow dynamics of the weights onto the inhibitory neuron) helps satisfying the condition and thus making the rule stable.

See Section 2.4 for a detailed analysis.

#### 1.6 *ForcedBalance* learning rule

The equations for the ForcedBalance learning rule are

$$\begin{aligned}\frac{dW_{EE}}{dt} &= +\alpha_1 g_E E(E_{set} - E) \\ \frac{dW_{EI}}{dt} &= \frac{1}{\tau_0}(W_{EIup} - W_{EI}) \\ \frac{dW_{IE}}{dt} &= +\alpha_3 g_I E(I_{set} - I) \\ \frac{dW_{II}}{dt} &= \frac{1}{\tau_0}(W_{IIup} - W_{II})\end{aligned} \tag{15}$$

and the conditions for the Up-state to be stable under this rule are

$$\begin{aligned}a_1 + b_1(W_{IIup}\, g_I + 1) < b_1'(W_{EEup}\, g_E - 1) \\ a_2 + b_2(W_{IIup}\, g_I + 1) < b_2'(W_{EEup}\, g_E - 1)\end{aligned} \tag{16}$$

where

$$\begin{aligned}a_1 &= (I_{set}\Theta_E\Theta_I\,\alpha_1\, g_E g_I + E_{set}^3\alpha_3)\, g_E g_I \\ b_1 &= I_{set}^2\Theta_E\,\alpha_1 g_E^2 g_I - E_{set}^2 I_{set}\,\alpha_1\, g_E^2 \\ b_1' &= E_{set}I_{set}\Theta_I\,\alpha_1\, g_E g_I^2 + E_{set}^2 I_{set}\,\alpha_3\, g_I^2 \\ a_2 &= 2\Theta_E\Theta_I\,\alpha_1\, g_E^2 g_I^2 \\ b_2 &= 2I_{set}\Theta_E\,\alpha_1\, g_E^2 g_I - E_{set}^2\,\alpha_1\, g_E^2 \\ b_2' &= 2E_{set}\Theta_I\,\alpha_1\, g_E g_I^2 + E_{set}^2\,\alpha_3\, g_I^2\end{aligned}$$

In Figure S5 we plot the stability condition of this rule, Eq. 16, for three different parameter values: $\alpha_1 = \alpha_3$, $\alpha_1 \gg \alpha_3$ (inhibitory plasticity slower); and $\alpha_1 \ll \alpha_3$ (excitatory plasticity slower).

In order to decide whether conditions Eq. 16 are satisfied in a more general case, note that $b_1$ and $b_2$ on the left-hand side are subtractions whereas $b_1'$ and $b_2'$ on the right-hand side are sums of positive definite terms, which helps satisfying the condition. On the other hand, one of the stability conditions of the neural subsystem might counter the effect: $(W_{IIup}\, g_I + 1)\tau_E > (W_{EEup}\, g_E - 1)\tau_I$ (see Section 2.2 below) but for biologically backed parameter values it is $\tau_E > \tau_I$ thus leaving room for the condition to hold. See Section 2.4 for a detailed analysis.

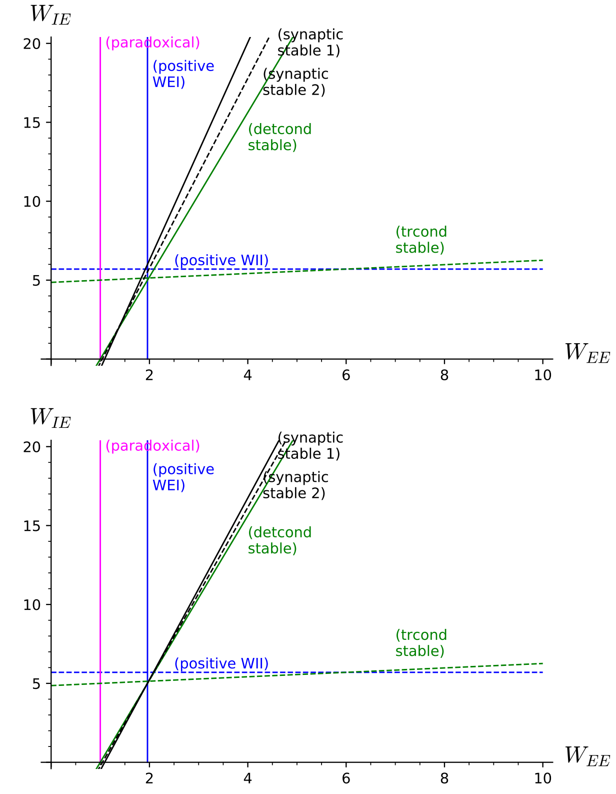

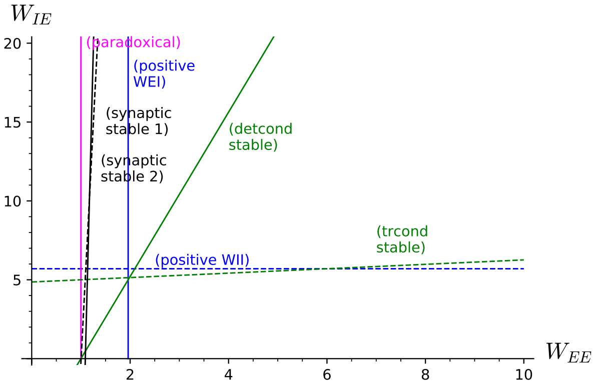

Figure S5: Regions of stability, *ForcedBalance* rule. (Top Left) Equal rates ($\alpha_{1,3} = 0.02$). (Top Right) Slow inhibitory ($\alpha_1 = 0.02$, $\alpha_3 = 0.0002$). (Bottom Left) Slow excitatory ($\alpha_1 = 0.0002$, $\alpha_3 = 0.02$). Every label is on the side where the corresponding condition holds (synaptic stable 1 and 2: Eq. 16 after switching to $W_{EE}$ and $W_{IE}$; detcond stable: Eq. 22; trcond stable: Eq. 23; positive $W_{EI}$: Eq. 25; positive $W_{II}$: Eq. 26; paradoxical: Eq. 27). Parameter values as in Table 1.

### 2 Detailed calculations

#### 2.1 Overview

We analyze the whole neural+synaptic system for every synaptic learning rule considered in this work, and study their stability. In every case, the general prescription is:

1. Take the combined neural+synaptic system and nondimensionalize all variables [see Sections 1.2 and 1.4 of Ref. 1][see Section 3.5 of Ref. 2], so that the two different time scales are evident (fast neural, slow synaptic).

2. Make a quasi-steady state (QSS) approximation of the neural subsystem [1, 2]. This means we will consider the neural subsystem is fast enough so that it converges "instantaneously" (when compared to the synaptic subsystem) to its corresponding fixed point. For this we will require that the stability conditions of the neural subsystem are satisfied (see below).

3. Find the steady-state solution of the synaptic subsystem, i.e. the Up-state fixed point; compute the Jacobian of the synaptic subsystem at the Up-state; compute the eigenvalues of the Jacobian [2, 3]. Two out of the four eigenvalues are expected to be zero because the Up-state is not an isolated fixed point of the system but a continuous 2D plane in 4D weight space.

4. Address (linear) stability. If both nonzero eigenvalues have negative real part, then the Up-state is stable under the learning rule; if at least one of the nonzero eigenvalues has positive real part, then the Up-state is unstable [2, 3]. (A note on abuse of notation: we might say indistinctly "the

Up-state is stable/unstable" and "the learning rule is stable/unstable".)

Eigenvalues and stability in the presence of continuous, i.e. non-isolated, attractors have been discussed in the context of neural networks for eye position control [4, 5] (keep in mind that the eigenvalues' critical value in these references is 1 instead of zero because they consider eigenvalues of the connectivity matrix alone, whereas we consider eigenvalues of the whole linear part). As the Up-state is a collection of non-isolated fixed points that form a 2D plane, there is no dynamics along the plane, and the linear stability analysis is enough to fully address stability—we do have two zero eigenvalues, but there is no need to compute the center manifold [3] because the other two eigenvalues represent the whole dynamics around the fixed point and have nonzero real part.

In order to apply the tools from Dynamical Systems' theory for flows in a unified way for both the neural and synaptic subsystems, we will switch from a discrete-time description of synaptic weight dynamics (where the change in weight $W$ is represented by $\Delta W$ applied every certain time interval) to a continuous-time description (where the weights are continuously evolving albeit with a long time scale $\tau_0$):

$$\Delta W \to \tau_0 \frac{dW}{dt}$$

In the following we first define the neural subsystem and compute its stability conditions (next subsection). Then we consider every learning rule in detail (following subsections).

**Paradoxical regime.** In this text we show detailed calculations of the stability conditions for the Homeostatic learning rule in the paradoxical regime only; see Section 2.5 for the non-paradoxical case.

#### 2.2 Neural dynamics

For the neural+synaptic system in the QSS approximation to be stable under a specific synaptic learning rule, it is necessary that the neural subsystem is stable so it remains in its QSS solution as the weights evolve. In this section we define the neural subsystem and compute its stability conditions.

(SageMath code in the Supplementary Material: `upstates-Neural subsystem stability.ipynb`)

##### 2.2.1 System's equations and fixed points

We consider a two-subpopulation model with firing-rate units $E$ and $I$ with ReLU activation functions (gain $g_X$, threshold $\Theta_X$, with $X = E, I$). The dynamics for synaptic currents above threshold is given by:

$$\begin{aligned} \frac{dE}{dt} &= \frac{1}{\tau_E}(-E + g_E(W_{EE}E - W_{EI}I - \Theta_E)) \\ \frac{dI}{dt} &= \frac{1}{\tau_I}(-I + g_I(W_{IE}E - W_{II}I - \Theta_I)) \end{aligned} \tag{17}$$

All variables and parameters are definite positive. In this subsection the synaptic weights $W_{XY}$ are fixed.

**Up-state.** The Up-state is the non-trivial fixed point of the system (i.e. a steady-state solution where $dE/dt = dI/dt = 0$):

$$\begin{aligned} E_{up} &= (W_{EI}\, g_I\, \Theta_I - (W_{II}\, g_I + 1)\, \Theta_E)\, g_E/C \\ I_{up} &= ((W_{EE}\, g_E - 1)\, \Theta_I - W_{IE}\, g_E\, \Theta_E)\, g_I/C \end{aligned} \tag{18}$$

where

$$C = W_{EI}W_{IE}\, g_E\, g_I - (W_{II}\, g_I + 1)(W_{EE}\, g_E - 1) \tag{19}$$

The activity of the excitatory and inhibitory subpopulations at the Up-state, $E_{up}$ and $I_{up}$, depend on all weight values. Only some of the combinations, however, lead to a stable steady state. We compute the stability conditions in the following subsection.

##### 2.2.2 Stability of neural fixed point (Up-state)

The Jacobian matrix, that is the matrix of first derivatives, gives information regarding the stability of fixed points: if the real parts of its eigenvalues are all negative, then the fixed point is stable.

The Jacobian of the neural system (Eq. 17) is

$$J = \begin{pmatrix} (W_{EE}g_E - 1)/\tau_E & -W_{EI}g_E/\tau_E \\ W_{IE}g_I/\tau_I & -(W_{II}g_I + 1)/\tau_I \end{pmatrix} \quad (20)$$

Its eigenvalues can be expressed as:

$$\lambda_{1,2} = \frac{1}{2}\left(Tr \pm \sqrt{Tr^2 - 4Det}\right) \quad (21)$$

where $Tr$ and $Det$ are the trace and determinant of the matrix, respectively. For eigenvalues either complex or purely real, their real parts are negative (and thus the Up-state is stable) when $Det > 0$ and $Tr < 0$, that is:

$$W_{EI}W_{IE}g_Eg_I > (W_{EE}g_E - 1)(W_{II}g_I + 1) \quad (22)$$

$$(W_{II}g_I + 1)\tau_E > (W_{EE}g_E - 1)\tau_I \quad (23)$$

Note that the positive determinant condition, Eq. 22, is equivalent to $C > 0$ (Eq. 19).

In the following, we will require that the stability conditions of the neural subsystem, Eqs. 22 and 23, are satisfied.

###### 2.2.3 Weight values consistent with an Up-state

The Up-state relationships, Eq. 18, are expressed as the $E$ and $I$ values resulting from a given set of weight values. If we set instead $E$ and $I$ to their target values $E_{set}$ and $I_{set}$ and solve for the weights, we get the weight values that are consistent with a given Up-state activity:

$$\begin{aligned} W_{EIup} &= \frac{(E_{set}W_{EEup} - \Theta_E)\,g_E - E_{set}}{I_{set}\,g_E} \\ W_{IIup} &= \frac{(E_{set}W_{IEup} - \Theta_I)\,g_I - I_{set}}{I_{set}\,g_I} \end{aligned} \quad (24)$$

Note first that any stable learning rule for the evolution of the weights for the neural subsystem (Eq. 17) must converge to weight values in accordance with these relationships (either in the form Eq. 24 or Eq. 18).

Second, note that the system is underdetermined and that is why two of the weights are free (chosen to be $W_{EE}$ and $W_{EI}$). Note also that all weight values must be positive; specifically, requiring $W_{EIup} > 0$ and $W_{IIup} > 0$ leads to

$$W_{EEup} > \frac{\Theta_E\,g_E + E_{set}}{E_{set}\,g_E} \quad (25)$$

$$W_{IEup} > \frac{\Theta_I\,g_I + I_{set}}{E_{set}\,g_I} \quad (26)$$

We refer to these expressions as the "positive $W_{EI}$" and the "positive $W_{II}$" conditions, respectively.

###### 2.2.4 Paradoxical effect

The paradoxical effect arises when an external depolarization of the inhibitory subpopulation (increase of $I$) produces an actual *decrease* of $I$. In this model, an external depolarization of $I$ can be mimicked by a decrease of its threshold $\Theta_I$, thus there is a paradoxical effect whenever the coefficient of $\Theta_I$ in the numerator of $I_{up}$ is positive. The coefficient is $g_I\,(W_{EE}\,g_E{-}1)/C$ and thus there is paradoxical effect if

$$W_{EE}\,g_E - 1 > 0 \quad (27)$$

The paradoxical effect can also be seen in a plot of the Up-state values $E_{up}$ and $I_{up}$ (Eq. 18) as a function of each individual weight. Specifically, from a naive point of view $I_{up}$ should increase when $W_{IE}$ is increased, and decrease when $W_{II}$ is increased; however, it does the opposite in either case (see Figure S6).

| | | | | | |
|---|---|---|---|---|---|
| $I_{set}$ | = | 14 | $E_{set}$ | = | 5 |
| $g_I$ | = | 4 | $g_E$ | = | 1 |
| $\Theta_I$ | = | 25 | $\Theta_E$ | = | 4.8 |
| $\tau_I$ | = | 2 | $\tau_E$ | = | 10 |

Table 1: Parameter values throughout the Supplementary Material. This set of parameter values makes the neural subsystem to be in the paradoxical regime (i.e. the Up-state is an inhibition-stabilized fixed point [6]). For non-paradoxical conditions, see Section 2.5.

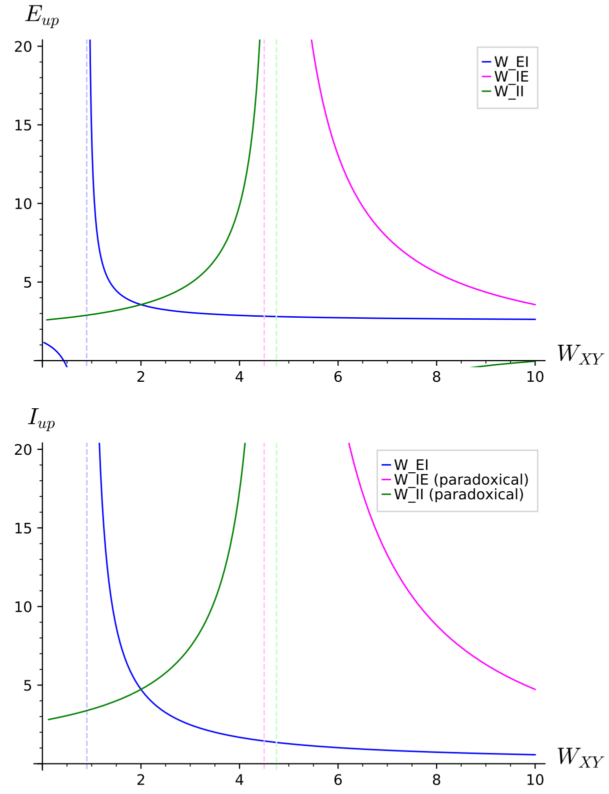

Figure S6: Paradoxical effect in the neural subsystem ($W_{EE} = 5$; parameter values as in Table 1). $E_{up}$ behaves as expected when each weight is varied. $I_{up}$, however, shows paradoxical behavior when either $W_{IE}$ or $W_{II}$ are varied. Dashed lines are the vertical asymptote of every case.

#### 2.3 *Homeostatic* learning rule: Detailed calculation

In this section we show in detail the calculation of the stability condition for the Homeostatic learning rule.

(SageMath code in the Supplementary Material: `upstates-Homeostatic stability.ipynb`)

##### 2.3.1 Definition of the learning rule

In continuous-time dynamics, the Homeostatic learning rule reads:

$$\begin{aligned} \frac{dW_{EE}}{dt} &= +\alpha_{EE}\, E(E_{set} - E) \\ \frac{dW_{EI}}{dt} &= -\alpha_{EI}\, I(E_{set} - E) \\ \frac{dW_{IE}}{dt} &= +\alpha_{IE}\, E(I_{set} - I) \\ \frac{dW_{II}}{dt} &= -\alpha_{II}\, I(I_{set} - I) \end{aligned} \tag{28}$$

where $\alpha_{XY}$ $(X, Y = E, I)$ are the learning rates (with appropriate units) setting the time scales of the weight dynamics, and $E_{set}$ and $I_{set}$ are the set points of the excitatory and inhibitory subpopulations, respectively.

The fixed points of the system (i.e. steady states) are determined by setting all derivatives to zero. There is a non-trivial fixed point compatible with the neural subsystem being above threshold: the Up-state, that is the set of weight values such that:

$$\begin{aligned} E_{up} &= E_{set} \\ I_{up} &= I_{set} \end{aligned} \tag{29}$$

The values of the weights corresponding to the Up-state are given by the (underdetermined) system defined by equating Eqs. 29 and 18. Since it is a two-equation system for a set of four unknown weights, there are two free weights that we choose to be $W_{EEup}$ and $W_{IEup}$. The values of the other two are given by Eq. 24. This means that the Up-state fixed point is actually a continuous set of non-isolated fixed points forming a 2D plane in 4D weight space. In other words, there is an infinite number of weight values compatible with the Up-state (possibly not all stable, though).

##### 2.3.2 Nondimensionalization

Next we nondimensionalize all variables in order to have a simpler system and make the QSS approximation in a safe way. We define new (nondimensional) variables $e$, $i$, $\tau$, $w_{EE}$, $w_{EI}$, $w_{IE}$, and $w_{II}$, and their corresponding scaling parameters. We substitute the new variables into the full system (neural+synaptic, Eqs. 17 and 28) and choose the values of the scaling parameters such that all nondimensional variables are of order 1 (see attached SageMath code). With this, the full system reads:

$$\begin{aligned}\epsilon_E \frac{de}{d\tau} &= -e + Rew_{EE} - \frac{iw_{EI}}{R} - \theta_E \\ \epsilon_I \frac{di}{d\tau} &= -i + \frac{Rew_{IE}}{g} - \frac{iw_{II}}{Rg} - \theta_I \\ \frac{dw_{EE}}{d\tau} &= -e(e-1) \\ \frac{dw_{EI}}{d\tau} &= +\alpha_2 i(e-1) \\ \frac{dw_{IE}}{d\tau} &= -\alpha_3 e(i-1) \\ \frac{dw_{II}}{d\tau} &= +\alpha_4 i(i-1)\end{aligned} \tag{30}$$

where we defined the new parameters

$$\begin{aligned}\epsilon_E &= \tau_E/\tau_0 \\ \epsilon_I &= \tau_I/\tau_0 \\ \tau_0 &= 1/(\alpha g_E E_{set} I_{set}) \\ R &= E_{set}/I_{set} \\ g &= g_E/g_I \\ \alpha_2 &= \alpha_{EI}/\alpha_{EE} \\ \alpha_3 &= \alpha_{IE}/\alpha_{EE} \\ \alpha_4 &= \alpha_{II}/\alpha_{EE} \\ \theta_E &= (g_E/E_{set})\Theta_E \\ \theta_I &= (g_I/I_{set})\Theta_I\end{aligned}$$

###### 2.3.3 Quasi-steady state approximation

Neural dynamics evolves in a much shorter time scale ($\tau_E$ and $\tau_I$) than synaptic dynamics ($\tau_0$):

$$\begin{aligned}\tau_E \ll \tau_0 &\implies \epsilon_E \ll 1 \\ \tau_I \ll \tau_0 &\implies \epsilon_E \ll 1\end{aligned}$$

which implies

$$\begin{aligned}\epsilon_E \frac{de}{d\tau} &\sim 0 \\ \epsilon_I \frac{di}{d\tau} &\sim 0\end{aligned} \tag{31}$$

thus we can safely assume $e$ and $i$ very quickly reach quasi-equilibrium values, i.e. practically instantaneous convergence to quasi-steady state (QSS) values as if the weights were fixed, while the synaptic weights evolve according to their slow dynamics. This allows us to reduce the system's dimensionality from six to four.

In the QSS approximation, the values of the nondimensionalized excitatory and inhibitory activities instantaneously track the slow dynamics of the learning rule. They are determined by applying Eq. 31 to the first two rows of Eq. 30; solving for $e$ and $i$ leads to

$$\begin{aligned}e_{qss} &= (g\theta_I w_{EI} - (w_{II} + Rg)\theta_E)/c \\ i_{qss} &= (Rg\theta_I(Rw_{EE} - 1) - R^2\theta_E w_{IE})/c\end{aligned} \tag{32}$$

where

$$c = Rw_{EI}w_{IE} - (w_{II} + Rg)(Rw_{EE} - 1)$$

The full system in the QSS approximation reads

$$\begin{aligned}\frac{dw_{EE}}{d\tau} &= -e_{qss}(e_{qss} - 1) \\ \frac{dw_{EI}}{d\tau} &= +\alpha_2 i_{qss}(e_{qss} - 1) \\ \frac{dw_{IE}}{d\tau} &= -\alpha_3 e_{qss}(i_{qss} - 1) \\ \frac{dw_{II}}{d\tau} &= +\alpha_4 i_{qss}(i_{qss} - 1)\end{aligned} \tag{33}$$

where $e_{qss}$ and $i_{qss}$ are nonlinear functions of the weights as defined by Eq. 32.

Note that the Up-state fixed point, defined by making all derivatives equal to zero, can be expressed as

$$\begin{aligned}e_{qss} &= 1 \\ i_{qss} &= 1\end{aligned} \tag{34}$$

which is the nondimensionalized version of Eq. 29. The weight values compatible with this condition are defined by equating Eqs. 32 and 34:

$$\begin{aligned}w_{EIup} &= R(Rw_{EEup} - 1) - R\theta_E \\ w_{IIup} &= R(Rw_{IEup} - g) - Rg\theta_I\end{aligned} \tag{35}$$

($w_{EEup}$ and $w_{IEup}$ are free). This is the nondimensionalized version of Eq. 24.

##### 2.3.4 Stability condition

The program for assessing linear stability of the Up-state is as follows: a) compute the Jacobian (the matrix of first derivatives) of Eq. 33 and evaluate it at the Up-state; b) compute the eigenvalues of the Jacobian (two of them will be zero because the fixed points form a continuous 2D plane in phase space); c) If the real part of the two nonzero eigenvalues is negative then the Up-state is stable; if at least one of the nonzero eigenvalue has positive real part then the Up-state is unstable.

**Jacobian matrix.** Let the full system in the QSS approximation (Eq. 33) be written as

$$\frac{dw_{EE}}{d\tau} = f_{EE}(e_{qss}, i_{qss})$$
$$\frac{dw_{EI}}{d\tau} = f_{EI}(e_{qss}, i_{qss})$$
$$\text{etc}\ldots$$

where $e_{qss}$ and $i_{qss}$ are functions of the weights as defined by Eq. 32. By applying the chain rule the elements $J_{ij}$ $(i, j = 1 \ldots 4)$ of the Jacobian matrix can be expressed as

$$J_{11} = \frac{df_{EE}}{dw_{EE}} = \frac{df_{EE}}{de_{qss}}\frac{de_{qss}}{dw_{EE}} + \frac{df_{EE}}{di_{qss}}\frac{di_{qss}}{dw_{EE}}$$
$$J_{12} = \frac{df_{EE}}{dw_{EI}} = \frac{df_{EE}}{de_{qss}}\frac{de_{qss}}{dw_{EI}} + \frac{df_{EE}}{di_{qss}}\frac{di_{qss}}{dw_{EI}}$$
$$J_{13} = \ldots$$
$$J_{21} = \frac{df_{EI}}{dw_{EE}} = \frac{df_{EI}}{de_{qss}}\frac{de_{qss}}{dw_{EE}} + \frac{df_{EI}}{di_{qss}}\frac{di_{qss}}{dw_{EE}}$$
$$J_{22} = \ldots$$
$$\text{etc}\ldots$$

In order to have the Jacobian specialized in the Up-state, these expressions are to be substituted by Eqs. 32-35.

**Eigenvalues of the Jacobian matrix.** The Jacobian matrix has two zero eigenvalues and two nonzero eigenvalues. The nonzero eigenvalues have the form:

$$\lambda_{\pm} = \frac{A \pm \sqrt{A^2 - DC}}{C} \tag{36}$$

where

$$\begin{aligned} A &= R^2 g\theta_I + (R^2\alpha_3 + \alpha_4)Rw_{EEup} \\ &\quad - (R^2 + \alpha_2)Rw_{IEup} + \alpha_2 g\theta_I - R^2\alpha_3 - \alpha_4 \\ C &= 2R(Rg\theta_I w_{EEup} - R\theta_E w_{IEup} - g\theta_I) \\ D &= 2(R^2\alpha_3 + \alpha_4)(R^2 + \alpha_2)/R \end{aligned} \tag{37}$$

**Sign of the eigenvalues.** To determine the sign of the real part of Eq. 36, first note that the factor $D$ is positive definite. Second, $C$ must be positive because it is related to one of the stability conditions of the neural subsystem (Eq. 22, after substituting back to dimensionalized quantities). Note next that $A^2 - DC$ is less than $A^2$ (since $C$ and $D$ are positive), and thus the square root is either real and less than $|A|$ or imaginary, both cases leading to $\text{Re}(A \pm \sqrt{A^2 - DC}) < 0$ if $A < 0$. The learning rule is then stable (both eigenvalues have negative real part) if $A < 0$, which in terms of the original parameters and free weights $W_{EE}$ and $W_{IE}$ reads:

$$\begin{aligned} &(E_{set}^2\alpha_{EE} + I_{set}^2\alpha_{EI})E_{set}g_E W_{IEup} \\ &\quad > + (E_{set}^2\alpha_{IE} + I_{set}^2\alpha_{II})I_{set}(W_{EEup}g_E - 1) \\ &\quad\quad + (E_{set}^2\alpha_{EE} + I_{set}^2\alpha_{EI})\Theta_I g_E \end{aligned} \tag{38}$$

##### 2.3.5 Analysis of the stability condition

It is hard to determine whether the stability condition Eq. 38 is satisfied for a general set of parameter values (see numerical analysis below). However, by using the Up-state relationship Eq. 24, this condition can be re-expressed in a more useful form in terms of $W_{EE}$ and $W_{II}$:

$$\begin{aligned} &(R^2\alpha_3 + \alpha_4)(W_{EEup}\, g_E - 1)g_I \\ &\quad < (R^2 + \alpha_2)(W_{IIup}\, g_I + 1)g_E \end{aligned} \tag{39}$$

Note that learning rates values of the same order lead to $\alpha_{2,3,4} \sim 1$ and that biologically backed parameter values satisfy:

$$I_{set} > E_{set}$$
$$g_I > g_E$$

both likely preventing the condition to hold.

On the other hand, small enough values of $\alpha_3$ and $\alpha_4$ (by making the dynamics of the weights onto the inhibitory neuron $W_{IE}$ and $W_{II}$ slower) would help satisfy the condition thus making the system stable.

###### 2.3.6 Relationship between the stability condition and the paradoxical condition

The boundary of the stability condition for this learning rule, Eq. 38, is a linear function in the $(W_{EE}, W_{IE})$ space with a slope that tends to infinity as the excitatory learning rates ($\alpha_{EE,EI}$) tend to zero:

$$\text{slope} = \frac{(E_{set}^2 \alpha_{IE} + I_{set}^2 \alpha_{II}) I_{set}}{(E_{set}^2 \alpha_{EE} + I_{set}^2 \alpha_{EI}) E_{set}}$$

while its root is a complicated expression (see SageMath notebook) that tends to $W_{EE} = 1/g_E$. The region of stability is to the left of the line. Thus, the boundary of stability in this limit coincides exactly with the boundary of the paradoxical condition ($W_{EE} > 1/g_E$). This can be construed as an inconsistency/contradiction between the stability of the rule and the existence of the paradoxical effect.

##### 2.4 Detailed calculations for the other rules

The stability calculations for the rest of the rules follow very similar paths. They can be found in the corresponding SageMath-Jupyter notebooks:

`upstates-CrossHomeostatic stability.ipynb`

`upstates-TwoTerm stability.ipynb`

`upstates-SynapticScaling stability.ipynb`

`upstates-ForcedBalance stability.ipynb`

##### 2.5 Stability of the rules in a non-paradoxical regime

All results above were developed with the neural subsystem set in the paradoxical regime—that is, the region in $(W_{EE}, W_{IE})$ leading to a stable Up-state was completely within the paradoxical region ($W_{EE} g_E > 1$). In order to show the importance of the paradoxical behavior for the stability of the learning rules, we also computed the stability conditions of every learning rule in a more general setting where the excitatory subpopulation in the neural subsystem has an external, constant, excitatory input current $I_{ext}$. This allows the neural subsystem to display both paradoxical and non-paradoxical stable behavior (in the second case, at the expense of the Up-state not being an inhibition-stabilized fixed point). The results can be found in the following SageMath-Jupyter notebooks:

`upstates-Neural subsystem stability-with Iext.ipynb`

`upstates-Homeostatic stability-with Iext.ipynb`

`upstates-TwoTerm stability-with Iext.ipynb`

`upstates-SynapticScaling stability-with Iext.ipynb`

`upstates-ForcedBalance stability-with Iext.ipynb`

(The stability condition of the Cross-Homeostatic learning rule doesn't depend on $I_{ext}$.)

#### 3 Learning rule from loss function

(SageMath code in the Supplementary Material: `upstates-Loss function.ipynb`)

Here we show how to compute the learning rule starting from a loss function. Then we make an approximation by considering the weight values are close to the values corresponding to an Up-state.

##### 3.1 General prescription

We consider the full neural+synaptic system in the QSS approximation (see e.g. Section 2.3). In this approximation the neural subsystem is represented by the quasi-steady-state values

$$\begin{aligned} E &= E_{up}(W_{EE}, W_{EI}, W_{IE}, W_{II}) \\ I &= I_{up}(W_{EE}, W_{EI}, W_{IE}, W_{II}) \end{aligned} \tag{40}$$

where the functions $E_{up}$ and $I_{up}$ are defined by Eq. 18 (see [7] for a related discussion on quasi-steady state, synaptic plasticity, and gradient descent).

The synaptic subsystem, that is the learning rule, will be obtained as a result of considering a specific loss function, and the general prescription to compute the learning rule from a loss function $L$ is the following:

1. Consider a loss function depending on $E$ and $I$ (which in turn depend on all weights):

   $$L = L(E, I)$$

   Conditions to be satisfied by the loss function are, for instance, to be smooth enough (i.e. continuous and differentiable) and to have a minimum when the activities $E$ and $I$ are at the set points $E_{set}$ and $I_{set}$ (i.e. homeostatic plasticity).

2. The dynamics of the weights is such that it follows a gradient descent on the loss function towards its minimum. In vector notation:

   $$\Delta \mathbf{W} = -\alpha \nabla L \tag{41}$$

   with a single learning rate $\alpha$ for simplicity. The unfolded learning rules, that is the equations that govern the weights' dynamics, are then

   $$\begin{aligned} \Delta W_{EE} &= -\alpha \frac{\partial L}{\partial W_{EE}} \\ \Delta W_{EI} &= -\alpha \frac{\partial L}{\partial W_{EI}} \\ \Delta W_{IE} &= -\alpha \frac{\partial L}{\partial W_{IE}} \\ \Delta W_{II} &= -\alpha \frac{\partial L}{\partial W_{II}} \end{aligned} \tag{42}$$

3. The partial derivatives of the loss function in Eq. 42 are:

   $$\begin{aligned} \frac{\partial L}{\partial W_{EE}} &= \frac{\partial L}{\partial E}\frac{\partial E}{\partial W_{EE}} + \frac{\partial L}{\partial I}\frac{\partial I}{\partial W_{EE}} \\ \frac{\partial L}{\partial W_{EI}} &= \frac{\partial L}{\partial E}\frac{\partial E}{\partial W_{EI}} + \frac{\partial L}{\partial I}\frac{\partial I}{\partial W_{EI}} \\ \frac{\partial L}{\partial W_{IE}} &= \frac{\partial L}{\partial E}\frac{\partial E}{\partial W_{IE}} + \frac{\partial L}{\partial I}\frac{\partial I}{\partial W_{IE}} \\ \frac{\partial L}{\partial W_{II}} &= \frac{\partial L}{\partial E}\frac{\partial E}{\partial W_{II}} + \frac{\partial L}{\partial I}\frac{\partial I}{\partial W_{II}} \end{aligned} \tag{43}$$

   or, in vector notation:

   $$\nabla L = \frac{\partial L}{\partial E}\nabla E + \frac{\partial L}{\partial I}\nabla I \tag{44}$$

   Here we use the chain rule for the derivatives because it gives us much more compact expressions at the end.

4. The partial derivatives in the gradients $\nabla E = \left(\frac{\partial E}{\partial W_{EE}}, \dots\right)$ and $\nabla I = \left(\frac{\partial I}{\partial W_{EE}}, \dots\right)$ etc. are to be taken from the quasi-steady-state values of $E$ and $I$, Eq. 40. We will, however, compute the partial derivatives from the implicit expressions given by setting $dE/dt = dI/dt = 0$ in Eq. 17 without solving for $E$ and $I$.

#### 3.2 Detailed calculation

##### 3.2.1 Exact learning rules

**Loss function.** We choose a very general loss function that depends homeostatically on both $E$ and $I$ activities:

$$L(E, I) = \frac{1}{2}(E_{set} - E)^2 + \frac{1}{2}(I_{set} - I)^2 \tag{45}$$

This loss function is an elliptic paraboloid in $(E, I)$ space with a global minimum at $(E_{set}, I_{set})$ so a gradient descend learning rule as above should converge to that minimum (see Liapunov function and gradient systems: [3, Section 1.1B][8, Sections 9.3 and 9.4][2, Section 7.2]). Keep in mind, however, that $L$ has a different shape when expressed as a function of the weights, and that $E$ and $I$ are not necessarily monotonic functions of the weights (particularly for a paradoxical system), so the conditions for the set point of $L$ to be stable or a global minimum or even unique are not necessarily satisfied.

**Partial derivatives of $L$.** The partial derivatives of $L$ with respect to $E$ and $I$ are simply

$$\begin{aligned}\frac{\partial L}{\partial E} &= -(E_{set} - E)\\ \frac{\partial L}{\partial I} &= -(I_{set} - I)\end{aligned} \tag{46}$$

**Partial derivatives of $E$ and $I$.** We compute the partial derivatives $\partial X/\partial W_{XY}$ $(X, Y = E, I)$ by first equating the neural subsystem (Eq. 17) to zero:

$$\begin{aligned}E &= g_E(W_{EE}E - W_{EI}I - \Theta_E))\\ I &= g_I(W_{IE}E - W_{II}I - \Theta_I))\end{aligned} \tag{47}$$

then differentiating the implicit functions:

$$\begin{aligned}\frac{\partial E}{\partial W_{EE}} &= g_E(E + W_{EE}\frac{\partial E}{\partial W_{EE}}) - g_E W_{EI}\frac{\partial I}{\partial W_{EE}}\\ \frac{\partial E}{\partial W_{EI}} &= g_E W_{EE}\frac{\partial E}{\partial W_{EI}} - g_E(I + W_{EI}\frac{\partial I}{\partial W_{EI}})\\ \frac{\partial E}{\partial W_{IE}} &= g_E W_{EE}\frac{\partial E}{\partial W_{IE}} - g_E W_{EI}\frac{\partial I}{\partial W_{IE}}\\ \frac{\partial E}{\partial W_{II}} &= g_E W_{EE}\frac{\partial E}{\partial W_{II}} - g_E W_{EI}\frac{\partial I}{\partial W_{II}}\\ \frac{\partial I}{\partial W_{EE}} &= g_I W_{IE}\frac{\partial E}{\partial W_{EE}} - g_I W_{II}\frac{\partial I}{\partial W_{EE}}\\ \frac{\partial I}{\partial W_{EI}} &= g_I W_{IE}\frac{\partial E}{\partial W_{EI}} - g_I W_{II}\frac{\partial I}{\partial W_{EI}}\\ \frac{\partial I}{\partial W_{IE}} &= g_I(E + W_{IE}\frac{\partial E}{\partial W_{IE}}) - g_I W_{II}\frac{\partial I}{\partial W_{IE}}\\ \frac{\partial I}{\partial W_{II}} &= g_I W_{IE}\frac{\partial E}{\partial W_{II}} - g_I(I + W_{II}\frac{\partial I}{\partial W_{II}})\end{aligned} \tag{48}$$

and then solving for the derivatives:

$$\begin{aligned}\frac{\partial E}{\partial W_{EE}} &= -(EW_{II}\, g_E\, g_I + Eg_E)/C\\ \frac{\partial E}{\partial W_{EI}} &= (IW_{II}\, g_E\, g_I + Ig_E)/C\\ \frac{\partial E}{\partial W_{IE}} &= EW_{EI}\, g_E\, g_I/C\\ \frac{\partial E}{\partial W_{II}} &= -IW_{EI}\, g_E\, g_I/C\\ \frac{\partial I}{\partial W_{EE}} &= -EW_{IE}\, g_E\, g_I/C\\ \frac{\partial I}{\partial W_{EI}} &= IW_{IE}\, g_E\, g_I/C\\ \frac{\partial I}{\partial W_{IE}} &= (EW_{EE}\, g_E - E)g_I/C\\ \frac{\partial I}{\partial W_{II}} &= -(IW_{EE}\, g_E - I)g_I/C\end{aligned} \tag{49}$$

where

$$C = W_{EI}W_{IE}\, g_E\, g_I - (W_{II}\, g_I + 1)(W_{EE}\, g_E - 1)$$

**Exact learning rules.** Putting everything together, the learning rules Eq. 42 are:

$$\begin{aligned}\Delta W_{EE} =& -\frac{\alpha}{C}((I_{set} - I)EW_{IE}\, g_e\, g_I\\ &+ (E_{set} - E)E(W_{II}\, g_I + 1)g_E)\\ \Delta W_{EI} =& +\frac{\alpha}{C}((I_{set} - I)IW_{IE}\, g_e\, g_I\\ &+ (E_{set} - E)I(W_{II}\, g_I + 1)g_E)\\ \Delta W_{IE} =& +\frac{\alpha}{C}((E_{set} - E)EW_{EI}\, g_e\, g_I\\ &+ (I_{set} - I)E(W_{EE}\, g_E - 1)g_I)\\ \Delta W_{II} =& -\frac{\alpha}{C}((E_{set} - E)IW_{EI}\, g_e\, g_I\\ &+ (I_{set} - I)I(W_{EE}\, g_E - 1)g_I)\end{aligned} \tag{50}$$

Note that these are very complicated, nonlinear expressions because both $E$ and $I$ depend on all weights via Eq. 47. Also the denominator $C$ depends on all weights (see previous paragraph).

##### 3.2.2 Approximation

We want simpler expressions for the learning rules. Note that the exact expressions above all have a homeostatic factor (either $E - E_{set}$ or $I - I_{set}$) and a presynaptic factor (either $E$ or $I$), while the rest are complicated expressions coming from the derivatives $\partial E/\partial W_{XY}$ and $\partial I/\partial W_{XY}$. We want to keep the homeostatic and presynaptic factors as they are while simplifying the rest of the expressions (explicit dependence on the weights including $C$) by performing a lowest-order Taylor series expansion of the explicit dependence of Eqs. 49 on the weights. Although this is not a textbook Taylor expansion of the full expressions, it is very informative because the results can be much more easily interpreted (for a similar approach see [7]).

We perform a zeroth-order approximation of the derivatives $\partial E/\partial W_{XY}$ and $\partial I/\partial W_{XY}$ as functions of the weights (i.e. while holding the presynaptic factors $E$ and $I$ constant) around the Up-state. In this approximation the weights are not small but close to their target values, represented by the relationships Eq. 24. By substituting the result in Eq. 43, we get the following approximated learning rules:

$$\begin{aligned}
\Delta W_{EE} &= +\alpha_E E(I_{set} - I) + \beta_E E(E_{set} - E) \\
\Delta W_{EI} &= -\alpha_E I(I_{set} - I) - \beta_E I(E_{set} - E) \\
\Delta W_{IE} &= -\alpha_I E(E_{set} - E) + \beta_I E(I_{set} - I) \\
\Delta W_{II} &= +\alpha_I I(E_{set} - E) - \beta_I I(I_{set} - I)
\end{aligned} \tag{51}$$

where

$$\begin{aligned}
\alpha_E &= \alpha g_E E_{set} W_{IEup}/D \\
\alpha_I &= \alpha A/D \\
\beta_E &= \alpha g_E B/D \\
\beta_I &= \alpha I_{set}(1 - W_{EEup} g_E)/D
\end{aligned}$$

and

$$\begin{aligned}
A &= E_{set} W_{EEup} g_E - \Theta_E g_E - E_{set} \\
B &= E_{set} W_{IEup} - \Theta_I \\
D &= \Theta_I W_{EEup} g_E - \Theta_E W_{IEup} g_E - \Theta_I
\end{aligned}$$

**Analysis.** Note that $\alpha_E$, $\alpha_I$, $\beta_E$, and $\beta_I$ are all constant. Furthermore, note that

- $A > 0$ as it is equal to the "positive $W_{EI}$" condition, Eq. 25;
- $B > 0$ as it is part of the "positive $W_{II}$" condition, Eq. 26;
- $D > 0$ as it is equal to the numerator of $I_{up}$, Eq. 18 (up to a positive factor), which must be positive because the denominator is.

Interestingly, note that the learning rate $\beta_I$ can be either negative or positive depending on whether the Up-state where the dynamics is converging to is paradoxical ($W_{EEup} g_E - 1 > 0$) or not ($W_{EEup} g_E - 1 < 0$). Note that the terms with $\alpha_{E,I}$ in the approximated learning rules, Eq. 51, are exactly equal to the Cross-Homeostatic rules, Eq. 6. Additionaly, the terms with $\beta_{E,I}$ are exactly equal to the Homeostatic rules, Eq. 28, unless $\beta_I < 0$ which would make the learning rule a Cross-Homeo-antiHomeo hybrid.
